# Body size-dependent energy storage causes Kleiber’s law scaling of the metabolic rate in planarians

**DOI:** 10.1101/332916

**Authors:** Jochen C. Rink, Albert Thommen, Steffen Werner, Olga Frank, Jenny Philipp, Oskar Knittelfelder, Yihui Quek, Karim Fahmy, Andrej Shevchenko, Benjamin M. Friedrich, Frank Jülicher

## Abstract

Kleiber’s law, or the ¾-power law scaling of the metabolic rate with body mass, is considered one of the few quantitative laws in biology, yet its physiological basis remains unknown. Here, we report Kleiber’s law scaling in the planarian *Schmidtea mediterranea*. Its reversible and life history-independent changes in adult body size over 2 orders of magnitude reveal that Kleiber’s law does not emerge from the size-dependent decrease in cellular metabolic rate, but from a size-dependent increase in mass per cell. Through a combination of experiment and theoretical analysis of the organismal energy balance, we further show that the mass allometry is caused by body size dependent energy storage. Our results reveal the physiological origins of Kleiber’s law in planarians and thus have general implications for understanding a fundamental scaling law in biology.

## Introduction

Body size varies strikingly across animal phylogeny. From small crustaceans weighing a few ng to Blue Whales weighing in excess of 150 000 kg, body mass variations span more than 16 orders of magnitude (Makarieva et al., 2008; Sears & Calambokidis, 2002). In spite of such tremendous variation in scale and physiology, the organismal metabolic rate (*P;* defined as the heat produced by the organism per unit time measured in Watts, which is related to the rate of oxygen consumption (McDonald, 2002)) nevertheless follows a general scaling relationship with body mass (*M*). As originally described by Kleiber in 1932 (Max Kleiber, 1932), *P* can be expressed by a power-law of the form *P = aM*^*b*^, with *b* being the scaling exponent and a proportionality constant *a*. Although reported values of *b* vary somewhat between studies or specific animal species, a value of *b* ≈ ¾ is typically observed (Banavar, Cooke, Rinaldo, & Maritan, 2014; Blaxter, 1989; Brody, 1945; Calder, 1984; Hemmingsen, 1960; M Kleiber, 1961; Peters, 1983; Schmidt-Nielsen, 1984; G B West & Brown, 2005; Whitfield, 2006) and this allometric relation between mass and metabolic rate is consequently referred to as the “three-quarter” or “Kleiber’s law”. This implies that the specific metabolic rate (*P*/*M*) decreases as body mass increases, which is commonly interpreted as reflecting a size-dependent decrease of cellular metabolic rates. Surprisingly, despite being known since more than 80 years and termed one of the few quantitative laws in biology (Geoffrey B West, 1999), the physiological basis of Kleiber’s law remains under intense debate. The fact that all animals, irrespective of physiology, habitat or life style, obey Kleiber’s law suggests a fundamental constraint in animal metabolism (G B West & Brown, 2005). Many hypotheses have been proposed that suggest a variety of origins of Kleiber’s law. A major class of hypotheses are based on internal physical constraints (Glazier, 2005), for example space-filling fractal transportation networks (Goeffrey B West, Brown, & Enquist, 1997) or size-dependent limitation of resource transport across external and internal body surfaces (Davison, 1955; Mcmahon, 1973). Another class of hypotheses concerns external ecological constraints, for example the optimization of body size for maximising reproductive fitness (Kozlowski & Weiner, 1997). However, experimental validations have proven difficult, due to inter-species differences in anatomy or ageing-associated physiological changes within a species. As a result, all hypotheses regarding the origins of Kleiber’s law remain controversial also because a suitable model system has not been established.

Flatworm laboratory models offer interesting opportunities in this respect. Although usually studied for their regenerative abilities and pluripotent adult stem cells (Reddien & Alvarado, 2004; Rink, 2012; Saló & Agata, 2012), the model species *S. mediterranea* and other planarians display tremendous changes in body size. They grow when fed and literally shrink when starving (Baguñà et al., 1990), which in *S. mediterranea* amounts to reversible body length fluctuations between ~0.5 mm and ~20 mm. Such >40-fold changes in body length in a laboratory model facilitate quantifications of *P* and *M* as pre-condition for applying and testing of theoretical approaches. Moreover, the commonly studied asexual strain of *S. mediterranea* and other asexual planarians do not seem to age, thus rendering their reversible size changes independent of organismal aging (Glazier, 2005). Previous studies of metabolic rate scaling in planarians suggest a size-dependence of O_2_-consumption (Daly & Matthews, 1982; Hyman, 1919), but the size dependence of *P* has so far not been systematically quantified. We here report that metabolic rate scaling in *S. mediterranea* indeed follows Kleiber’s law and we apply a combination of experiments and theory to understand its physiological basis. Our analysis of the organismal energy balance reveals that the size-dependent decrease in the specific metabolic rate does not reflect a decrease in the metabolic rate per cell, but instead an increase in the mass per cell. Further, we demonstrate that the cell mass allometry reflects a size-dependent increase in lipid and glycogen stores. Our results therefore demonstrate that size-dependent energy storage causes Kleiber’s law scaling in planarians.

## Results

### Planarians display Kleiber’s law scaling of the metabolic rate

Kleiber’s law describes the scaling of metabolic rate with the mass of animals. In order to test whether the tremendous body size fluctuations of *S. mediterranea* (*Figure 1A*) follow Kleiber’s law, we needed to devise methods to accurately quantify the mass and metabolic rate of planarians.

**Figure 1.**
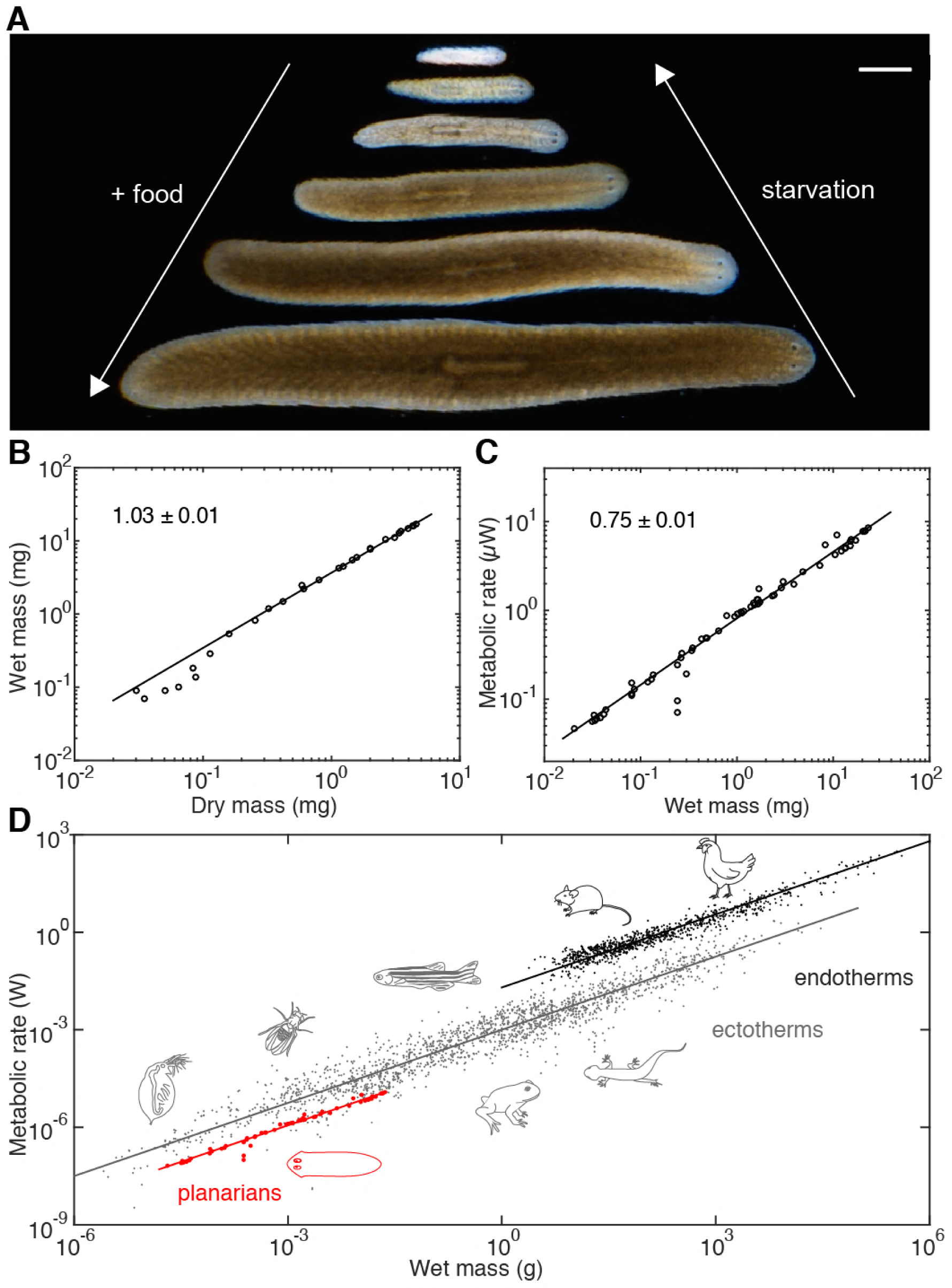
Kleiber’s law scaling during *S. mediterranea* body size changes. (**A**) Feeding (growth) and starvation (degrowth) dependent body size changes of *Schmidtea mediterranea*. Scale bar, 1 mm. (**B**) Wet versus dry mass scaling with body size. The scaling exponent ± standard error was derived from a linear fit for wet mass > 0.5 mg and represents the exponent *b* of the power law *y = ax*^*b*^. See *Figure 1 – source data 1* for numerical data. (**C**) Metabolic rate versus wet mass scaling by microcalorimetry. The metabolic rate was determined by a horizontal line fitted to the stabilised post-equilibration heat flow trace (*Figure 1 – figure supplement 1*) and the post-experimental dry mass determination of all animals in the vial was re-converted into wet mass by the scaling relation from (B). Each data point represents a vial average of a size-matched cohort. The scaling exponent ± standard error was derived from linear fits and represents the exponent *b* of the power law *y = ax*^*b*^. (**D**) Metabolic rate versus wet mass scaling in planarians from (C) (green) in comparison with published interspecies comparisons (Makarieva et al., 2008) amongst ectotherms (grey) or endotherms (black). Dots correspond to individual measurements; black and blue solid lines trace the ¾ scaling exponent; red line, linear fit to the planarian data. By convention (Makarieva et al., 2008), measurements from homeotherms obtained at different temperatures were converted to 37 °C, measurements from poikilotherms and our planarian measurements to 25 °, using the following factor: 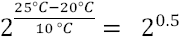 (20 °C: planarian data acquisition temperature).

To measure mass, we quantified both the dry and wet mass of individual planarians. Though dry mass measurements avoid the challenging removal of residual water from the mucus-coated animals, they are lethal and can therefore only be carried out once. As shown in *Figure 1B*, the wet and dry mass of *S. mediterranea* vary over > 2 orders of magnitude. Moreover, the near-constant ratio between wet and dry mass (~ 5; implying 80% water content) indicates minimal variations of the water content and thus facile interconversion of the two mass measurements.

In order to quantify metabolic rate, we measured the heat generated by live planarians of different sizes via microcalorimetry. Microcalorimetry measures the integrated heat generated by metabolic processes inside the animal and therefore provides a pathway-independent measure of total metabolic activity (Kemp & Guan, 1997). Cohorts of equally-sized 2 week-starved animals were enclosed in vials and their heat emission measured over a period of > 24 h (*Figure 1 – figure supplement 1*). Since animals were not immobilized, our measurements effectively reflect the routine metabolic rate that is generally used for aquatic animals (Dall, 1986). As shown in *Figure 1C*, the metabolic rate measurements increase with mass over nearly 3 orders of magnitude (from ~ 0.02 to 10 µW). The data points can be fit with a single power law that accurately describes the size-dependence of the metabolic rate across the entire size range. Intriguingly, the value of the scale exponent is 0.75 ± 0.01 and thus identical with the ~ 0.75 exponent associated with Kleiber’s law in inter-species comparisons. Consequently, the slope of the planarian data points (red) exactly parallels the characteristic slope of extensive published data sets of specific metabolic rate measurements (Makarieva et al., 2008) (*Figure 1D*). While the offsets between endo-and ectotherm traces might reflect different temperature regimes as previously noted (Hemmingsen, 1960; Makarieva et al., 2008), the common slopes stresses the universal nature of the ¾ law exponent across animal phylogeny. The fact that the same power law exponent is associated with the entire growth/degrowth-dependent body size interval of a planarian suggests that the same underlying principles are at work and that *S. mediterranea* is therefore a suitable model system for probing the physiological basis of Kleiber’s law.

### Size-dependence of planarian growth/degrowth dynamics

The physiological causes of planarian body size fluctuations are growth and degrowth. Therefore, understanding their underlying regulation might provide insights into the size-dependence of the metabolic rate. Planarian body size measurements are challenging due to their soft and highly deformable bodies. We therefore adapted our semi-automated live-imaging pipeline that extracts size measurements from multiple movie frames displaying the animals in an extended body posture (Werner, Rink, Riedel-Kruse, & Friedrich, 2014). We found that plan area provides the most robust size measure (*Figure 2 – figure supplement 1* and (Werner et al., 2014)), which we therefore use in the following. One first important question is to what extent size changes reflect a change in cell number. Since previous cell number estimates produced conflicting results (Romero & Baguñà, 1991; Takeda, Nishimura, & Agata, 2009) we developed two independent assays. First, we combined cell dissociation (Romero & Baguñà, 1991) with automated counting of fluorescently stained nuclei (*Figure 2A, top* and *Figure 2 – figure supplement 2*). Second, we used quantitative Western blotting to quantify the amount of the core Histone H3 in whole worm lysates, which we found to increase linearly with the number of cells (*Figure 2A, bottom*). Applying both assays to individually sized *S. mediterranea* revealed a close agreement between the two methods and scaling of cell numbers with plan area by a power law with the exponent 1.19 (*Figure 2B*), consistent with the previous conclusion that planarian body size changes largely reflect changes in cell numbers (Baguñà et al., 1990). Further, knowledge of the cell number/area scaling law allows the accurate interconversion of plan area into cell numbers in the experiments below.

**Figure 2.**
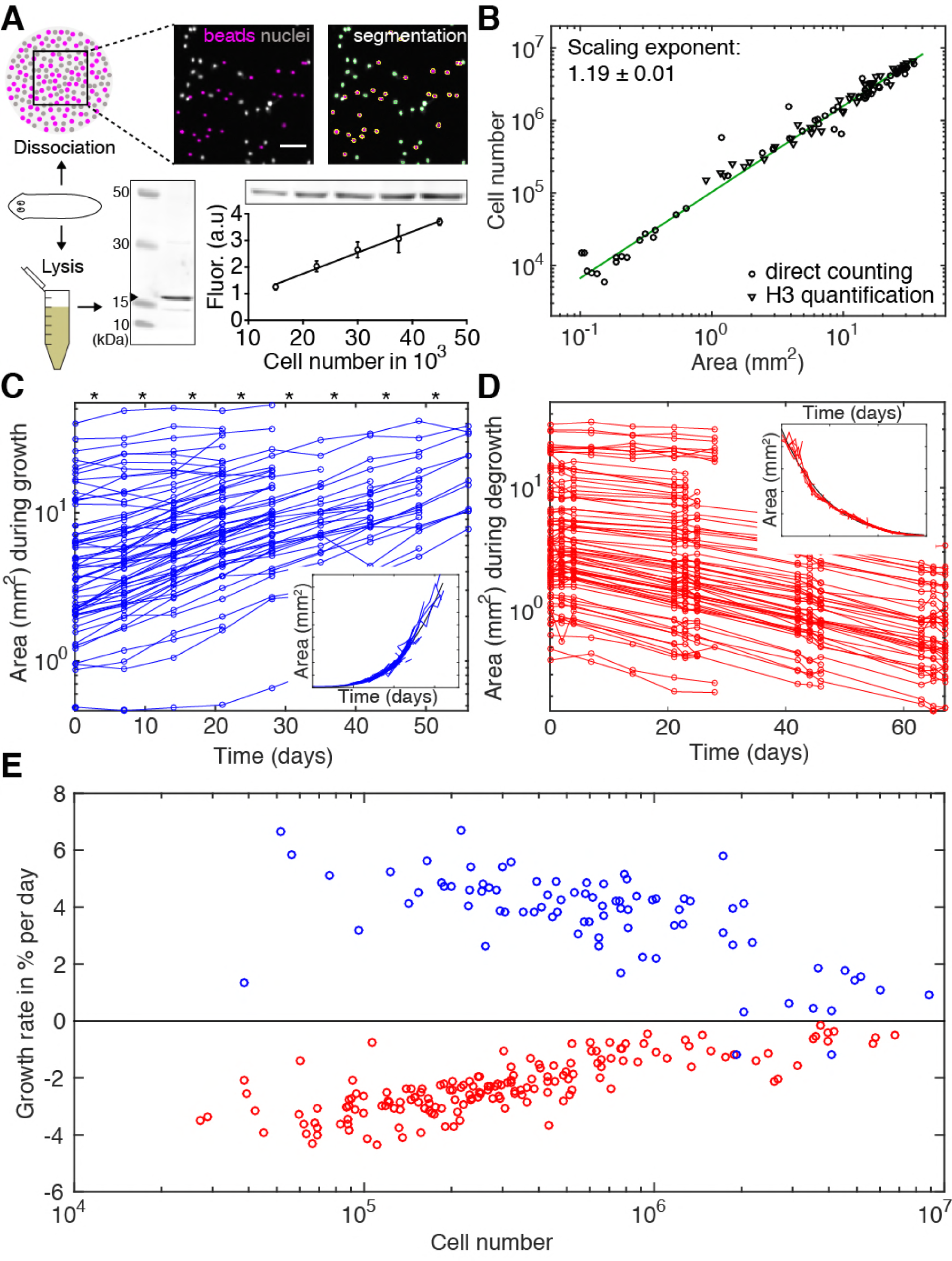
Growth and degrowth dynamics in *S. mediterranea*. (**A**) Assays to measure organismal cell numbers. (Top) image-based quantification of nuclei (grey) versus tracer beads (magenta) following whole animal dissociation in presence of the volume tracer beads. Bottom, Histone H3 protein quantification by quantitative Western blotting, which scales linearly with the number of FACS-sorted cells (bottom right). The line represents a fitted linear regression (data of 4 technical replicates) and serves as standard for calibrating the H3 band in planarian lysates (bottom left) run on the same gel into cell numbers. Values are shown as mean ± standard deviation. (**B**) Organismal cell number versus plan area scaling, by nuclei counts (circles) or Histone H3 protein amounts (triangles) (see also *Figure 2 – supplement 1 and 2)*. The scaling exponent ± standard error was derived from a linear fit and represents the exponent *b* of the power law *y = ax*^*b*^. Each data point represents one individual animal and the mean of several technical replicates, Histone H3 method: 9 independent experiments including 5 animals each; image-based approach: 4 independent experiments including 18, 10, 10 and 12 animals each. See *Figure 2 – source data 1-4* for numerical data. (**C**) Plan area changes of individual animals during growth. * indicate feeding time points (1x per week). Inset, concatenation of individual growth traces by area overlap. (**D**) Plan area change of individual animals during degrowth. Inset, concatenation of individual degrowth traces by area overlap. (**E**) Size-dependence of growth (blue) and degrowth rates (red) (see also *Figure 2 – figure supplement 3*). Individual data points were calculated by exponential fits to traces in (C) and (D) (growth: 2 overlapping time windows, degrowth: 3 overlapping time windows) and using the cell number/area scaling law from (B) to express rates as % change in cell number/day. The positive growth rates and negative degrowth rates are plotted on the same axis to facilitate comparison of size dependence. See *Figure 2 – source data 4* for data of (C) and (D).

To measure growth and degrowth rates, we quantified the change in plan area of individual *S. mediterranea* subjected to feeding at regular time intervals (*Figure 2C*) or continuous starvation (*Figure 2D*). Although individual measurements were noisy due to the aforementioned size quantification challenges, the data on >100 animals cumulatively reveal stereotypic time trajectories for both growth and degrowth (*Figure 2C-D, insets*). Therefore, planarian growth and degrowth dynamics are highly coordinated at the organismal level and our data are suitable for extracting the underlying rate constants. As shown in *Figure 2E*, we found that the growth rate of *S. mediterranea* decreases with body size, consistent with previous data (Baguñà et al., 1990). Unexpectedly, our analysis additionally revealed a similar size dependence of the degrowth rate. Interestingly, both growth and degrowth rates appeared to be largely independent of feeding history and thus solely a function of size (*Figure 2 – figure supplement 3A*). Taken together, our findings demonstrate that not only the specific metabolic rate (*Figure 1C-D*), but also the growth/degrowth rates decrease with body size in *S. mediterranea*.

### Systems-level control of planarian growth dynamics

Since growth reflects the metabolic assimilation of environmental resources and degrowth their subsequent catabolism, both are related to the overall metabolic rate of the animal. Therefore, the size dependence of growth/degrowth (*Figure 2E*) and metabolic rate (*Figure 1C-D*) might reflect a common physiological origin of the underlying scaling laws. We therefore devised a theoretical framework of planarian growth/degrowth as a function of the metabolic energy budget (*Figure 3A*). The central element of our model and previous approaches (Hou et al., 2008; Kooijman, 2009) is the organismal energy content *E*, which represents the sum of all physiologically accessible energy stores (e.g., carbohydrates, lipids and proteins). The energy content *E* fuels all metabolic processes within the animal, which collectively convert *E* into heat as measured by our microcalorimetry approach (*Figure 1C-D*). Hence, starvation results in a decrease of *E*, overall net catabolism and degrowth. However, *E* increases if the influx of energy obtained from the food (*J*) exceeds the energy lost through heat (*P*), which leads to net assimilation of resources and thus growth. The fact that planarians grow/degrow largely by a change in total cell numbers (*Figure 2B*) (Baguñà et al., 1990; Romero & Baguñà, 1991), fundamentally interconnects the organismal energy balance to organismal cell numbers. While excess energy from food intake stimulates increased cell proliferation (Baguñà, 1974) and growth, a starvation-induced net loss of energy decreases total cell numbers and thus, body size. Therefore, our framework relates changes in cell number during growth/degrowth to the energy content of the animal (*Figure 3A*). Importantly, our model does not make any assumptions regarding the underlying cellular or metabolic mechanisms, but simply states the physical energy balance of planarians.

**Figure 3.**
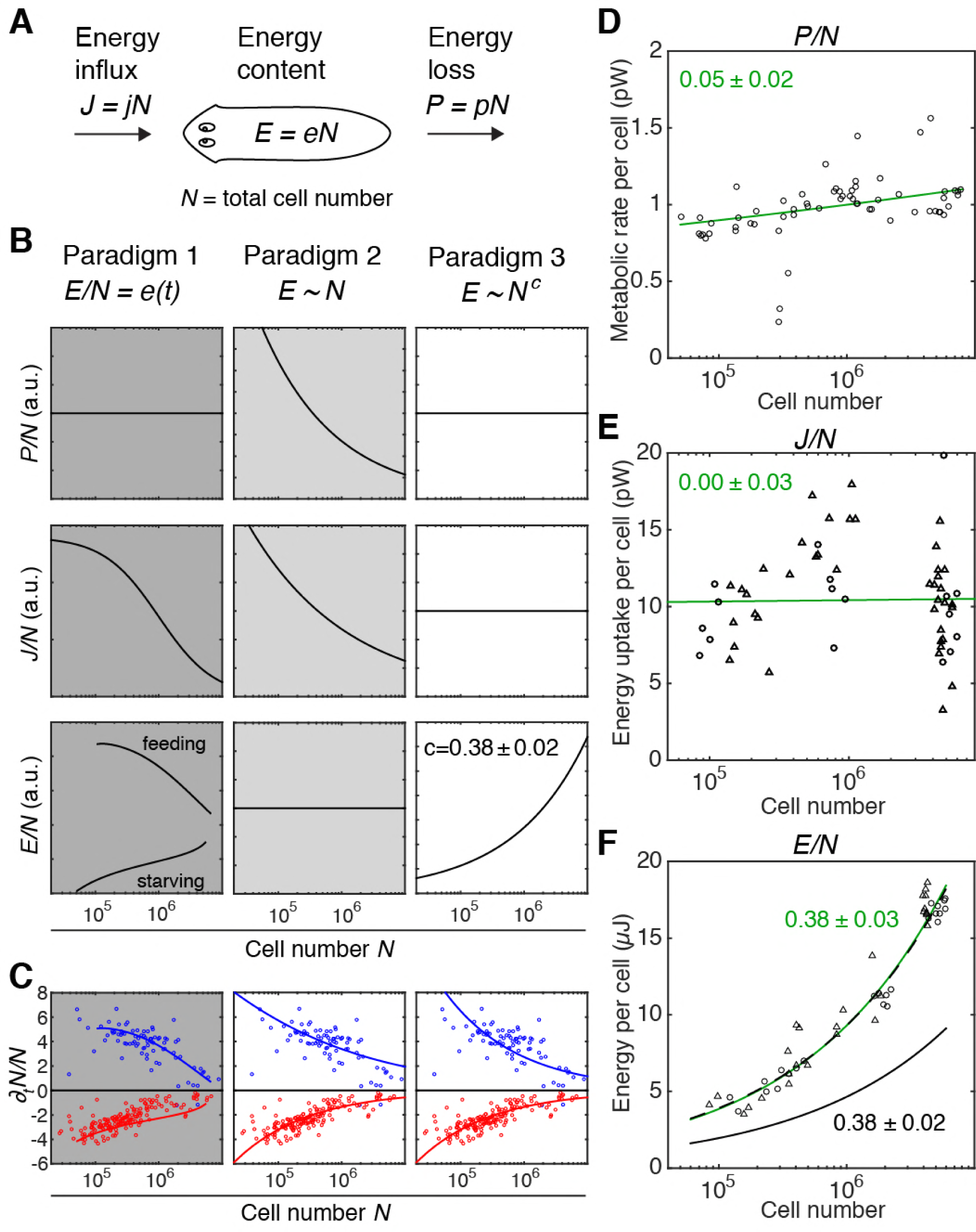
Size-dependent scaling of energy content explains growth/degrowth dynamics. (**A**) Planarian energy balance model. At the organismal level, changes in the physiological energy content *E* result from a change in the net energy influx *J* (feeding) and/or heat loss *P* (metabolic rate). Dividing *E, J* and *P* by the total cell number *N* approximates the energy balance on a per-cell basis. (**B**) Three hypothetical control paradigms of *E* during growth and degrowth (columns), which make specific predictions regarding the size-dependence of *J/N, E/N* and *P/N* (rows). Prediction traces and scale exponents were generated by modelling the measured growth/degrowth rates (*Figure 2E*) with the indicated control paradigm assumptions (see also *Figure 3 – figure supplement 1-3*). (**C**) Fit of the three control paradigms to the measured growth/degrowth rates (Fig. 2E). (**D**) Metabolic rate per cell (*P/N*) versus organismal cell number (*N*). Data points were derived by conversion of the measurements from the metabolic rate*/*dry mass scaling law (*Figure 1 – figure supplement 1B*) via the measured cell number/plan area (*Figure 2B*) and plan area/dry mass conversion laws (*Figure 3 – figure supplement 3A*). The scaling exponent ± standard error was derived from the respective linear fits (green line) and represents the exponent *b* of the power law *y = ax*^*b*^. (**E**) Energy uptake per cell versus organismal cell number (*N*). Data points reflect single-animal quantifications of ingested liver volume per plan area as shown in *Figure 3 – figure supplement 3D*, converted into energy intake/cell using the plan area/cell number scaling law (Fig. 2B) and the assumption that 1 µl of liver paste correspond to 6.15 J (“Nutrient report of calf liver,” 2016; Overmoyer, McLaren, & Brittenham, 1987). Circles, 2 weeks-starved and triangles, 3 weeks-starved animals. The scaling exponent ± standard error was derived from linear fits (green line) and represents the exponent b of the power law *y = ax*^*b*^. (**F**) Energy content per cell (*E/N*) versus organismal cell number (*N*). Data points reflect bomb calorimetry quantifications of heat release upon complete combustion of size matched cohorts of known dry mass as shown in *Figure 3 – figure supplement 3E*, converted via the measured cell number/plan area (Figure 2B) and plan area/dry mass conversion laws (*Figure 3 – figure supplement 3A*). Circles, 1 week-starved and triangles, 3 weeks-starved animals. The scaling exponent ± standard error was derived from linear fits (green line) to the data and represents the exponent *b* of the power law *y = ax*^*b*^. Solid black line, prediction from model 3 for the physiological energy content per cell assuming a constant metabolic rate *P/N* = 1 pW. Dashed line corresponds to respective prediction under the assumption that the physiological energy (solid black line) amounts to 50 % of combustible gross energy in the animal. See *Figure 3 – source data 1* for numerical data of (C)-(F).

With our quantitative data as experimental constraint (*Figure 2C-E*), the model allows us to explore hypothetical systems-level control paradigms of growth/degrowth dynamics (see also *Figure 3 – figure supplement 1*). In the first paradigm (*Figure 3B, left column*), dynamic changes in the organismal energy content depend on feeding conditions. Changes in cell number (e.g., rates of cell division and/or cell death) depend on the energy content per cell (*Figure 3 – figure supplement 2*). Consequently, two planarians with the same cell number might have different energy levels depending on the respective feeding history. In paradigm 2 (*Figure 3B, centre column*), the energy content is proportional to total cell number, i.e. it scales isometrically. Thus, growth occurs when “surplus” energy obtained from food is converted into new cells, whereas degrowth is the consequence of catabolism of existing cells in order to replenish metabolic energy. In paradigm 3, the energy content is also tightly coupled to cell number, but scales in a size-dependent manner with a characteristic exponent *c*, i.e. it scales allometrically. Theoretical analysis reveals that all three paradigms can approximate the measured growth/degrowth dynamics (*Figure 3C*). However, they differ in their specific predictions of the scaling behaviours of *E, P* and *J* with organismal cell number *N* (*Figure 3B*).

In order to experimentally distinguish between the paradigms, we consequently quantified *E, P* and *J* as a function of cell number (*N*). In order to obtain values for *P*/*N* (metabolic rate/cell), we converted our measurements of *P* as a function of dry mass (*Figure 1 – figure supplement 1B*) using the scaling laws for *N* and dry mass with plan area (*Figure 2B* and *Figure 3 – figure supplement 3A*). As shown in *Figure 3D*, the *P*/*N* estimates are of the order of 1 pW, similar to the average metabolic rate of a human cell (Bianconi et al., 2013; Purves & Sadava, 2004). Further, *P*/*N* is essentially independent of organismal cell number and animal size (scale exponent 0.05 +/-0.02), which rules out paradigm 2 (*Figure 3B*) as possible control principle. The size *independence* of *P*/*N* is further intriguing, as it implies that the size *dependence* of *P/M* as foundational basis of Kleiber’s law originates from size dependencies of *M/N* (mass per cell; see below).

To measure the food intake *J*, we developed an assay based on the homogenous dispersion of a known amount of small fluorescent beads in a known volume of planarian sustenance food (liver paste). Lysis of pre-sized animals immediately after feeding and quantification of bead numbers in the lysate thus provided a measure of the ingested food volume as a function of size (*Figure 3 – figure supplement 3B-D*). Although individual measurements varied significantly (likely reflecting inter-animal differences under our *ad libitum* feeding conditions), *J*/*N* did not display a clear size dependence (exponent 0.00 ± 0.03) (*Figure 3E*). Therefore, the volume of ingested food and thus energy uptake remains proportional to organismal cell number across the entire size range, which argues against both paradigms 1 and 2 (*Figure 3B*).

To approximate the energy content *E* of entire worms, we turned to bomb calorimetry. This method quantifies the heat release upon complete combustion of dried tissue in pure oxygen, thus providing a measure of gross energy content (McDonald, 2002). Our assay conditions allowed reproducible quantification of *E* of as little as 3 mg of dried tissue (*Figure 3 – figure supplement 3E*), corresponding to 200 planarians of a length of 2 mm (*Figure 3 – figure supplement 3A* and *Figure 2 – figure supplement 1D*). Intriguingly, *E*/*N* significantly increased with organismal cell numbers (scaling exponent 0.38 ± 0.03, *Figure 3F*), as assumed by paradigm 3 (*Figure 3B*). Moreover, the experimentally measured scaling exponent of the energy content agrees quantitatively with the prediction of paradigm 3 on basis of the experimentally measured growth/degrowth rates (*Figure 3F*; black solid line). The experimentally measured gross energy content and the physiologically accessible energy content *E* (green and black solid lines in *Figure 3F*) differ by a constant factor (2). The fact that the scaling exponent follows the prediction of paradigm 3 demonstrates the quantitative agreement between model and experiment and identifies size-dependent energy storage as systems-level control paradigm of planarian growth/degrowth dynamics.

### Size dependence of physiological energy stores

Since animals store energy in the form of biochemical compounds, size-dependent energy storage should consequently result in biochemically measurable effects. Little is currently known about planarian energy metabolism, but animals generally store metabolic energy in the form of triglycerides (TGs) inside lipid droplets (Birsoy, Festuccia, & Laplante, 2013). We therefore stained cross-sections of large and small animals with the lipid droplet marker LD540 (Spandl, White, Peychl, & Thiele, 2009). Both revealed prominent lipid droplets primarily within the intestinal epithelium, thus suggesting that the planarian intestine serves as a fat storage organ, as in *C. elegans* (Mak, 2012). However, the amount and size of the droplets per cell notably increased in large animals (*Figure 4A*). To obtain a quantitative measure of the lipid content size-dependence, we optimized total lipid extraction for planarians (*Figure 4 – figure supplement 1A*) and used mass spectrometry to measure the absolute amounts of various lipid classes (*Figure 4 – figure supplement 1B*). The 88-fold increase in TGs per unit cell in large planarians as compared to small animals (*Figure 4B*) demonstrates a striking size dependence of lipid stores in *S. mediterranea*.

**Figure 4.**
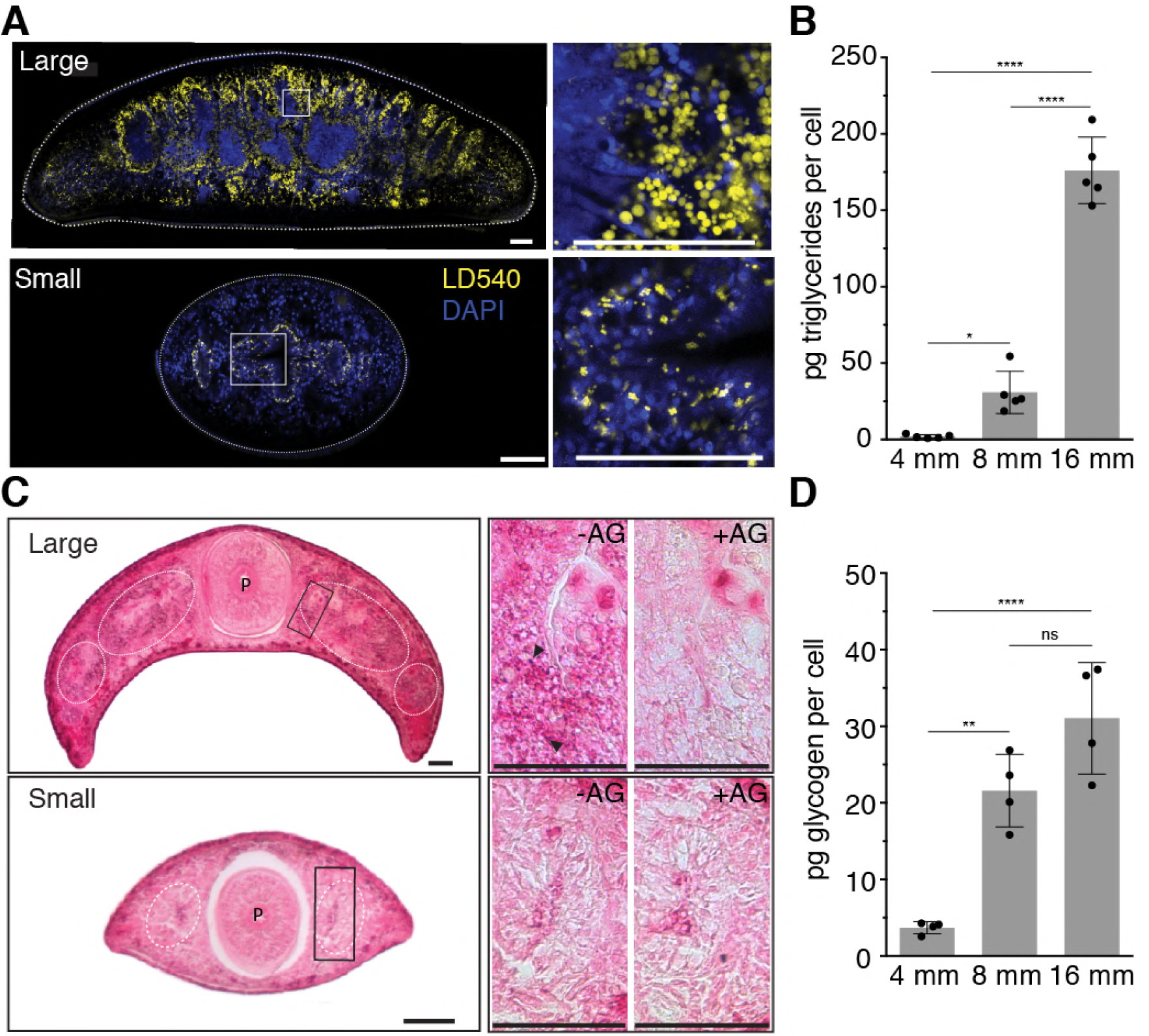
Size-dependence of lipid and glycogen storage. (**A**) Lipid droplet (LD540, yellow) (Spandl et al., 2009) and nuclei (DAPI, blue) staining of pre-pharyngeal transverse cross sections of a large (16 mm length, top left) and a small (4 mm, bottom left). Right, magnified view of the boxed areas to the left. Scale bars, 100 µm. See *Figure 4 – source data 1* for raw images. (**B**) Mass-spectrometry-based quantification of triglycerides in animals of the indicated size (*Figure 4 – figure supplement 1A-B*). All values were normalised to organismal cell numbers using the previously established length versus area (*Figure 2 – figure supplement 1D*) and *N*/*A* (*Figure 2B*) scaling laws. Bars mark mean ± standard deviation. n=5 biological replicates consisting of 40 pooled 4 mm, 20 8 mm and 6 16 mm long animals analysed in 2 technical replicates. Significance assessed by one-way ANOVA, followed by Tukey’s post-hoc test (\**p*_*adj*_≤0.05, \*\*\*\**p*_*adj*_≤0.0001). See *Figure 4 – source data 2* for numerical data and statistics. (**C**) Histological glycogen staining (Best’s Carmine method) of pre-pharyngeal transverse cross sections of a large (16 mm, top left) and a small (4 mm, bottom left). White circles: outline of intestine branches. P: Pharynx. Right, magnified view of the boxed areas to the left (black rectangles). +AG, pre-treatment with amyloglucosidase, which degrades glycogen; - AG, no pre-treatment of adjacent section. Arrow heads point to small, densely staining glycogen granules. Scale bars, 100 µm. See *Figure 4 – source data 1* for raw images. (**D**) Quantification of organismal glycogen content using an enzyme-based colorimetric assay in animals of the indicated length (*Figure 4 – figure supplement C-F*). Bars mark mean ± standard deviation. n=4 biological replicates (independent experiments), 40 pooled 4 mm, 20 8 mm, 8 16 mm analysed in 3 technical replicates. Significance assessed by one-way ANOVA, followed by Tukey’s post-hoc test (ns not significant, \*\**p*_*adj*_≤0.01, \*\*\*\**p*_*adj*_≤0.0001). See *Figure 4 – source data 2* for numerical data and statistics.

To further assess a possible size dependence of carbohydrate stores, we applied Best’s Carmine stain to cross-sections of large and small animals in order to probe for glycogen granules (*Figure 4C, left*). Comparison between adjacent sections with and without amyloglucosidase pre-treatment as specificity control (*Figure 4C, right*) together with the expression patterns of glycogen synthesis genes (*Figure 4 – figure supplement 1C*) both indicate a storage role of the planarian intestine for glycogen granules, thus again emphasizing the organ’s likely central role in energy homeostasis. Interestingly, also the intensity of glycogen staining appeared stronger in large animals (*Figure 4C, right*) and the quantification of glycogen content in animal extracts by an enzyme-based assay (*Figure 4 – figure supplement 1D-F*) demonstrated a > 8-fold increase in the amount of glycogen/cell in large over small animals (*Figure 4*). Therefore, both the lipid and carbohydrates stores are strongly size-dependent in *S. mediterranea*, which conclusively confirms our model’s prediction of size-dependent energy storage as a systems-level control paradigm of planarian growth and degrowth.

### Energy reserves and cell number govern Kleiber’s law in planarians

The size-dependent increase in the mass of lipid and glycogen stores is intriguing also in light of the previous indications that Kleiber’s law in planarians might originate from a size-dependent increase in mass per cell, rather than a decrease in metabolic rate (*Figure 3D*). To explore this potential link between the regulation of growth dynamics and Kleiber’s law, we first investigated the relative contributions of mass allometries to the emergence of the ¾ exponent. As a direct test, we derived the size dependence of cell numbers versus mass, using the various scaling laws established during the course of this study. As shown in *Figure 5A*, cell numbers scale with wet and dry mass with scale exponents of 0.74 ± 0.01 and 0.72 ± 0.01, respectively. This demonstrates that the mass per cell indeed increases disproportionately with size and with a very similar scaling exponent as for Kleiber’s law (*Figure 1C*). In conjunction with the practically size-independent scaling of cell number and metabolic rate (Figure 5B, scaling exponent 0.96 ± 0.02), these data demonstrate conclusively that the ¾ exponent of the metabolic rate/mass scaling law derives from the underlying scaling law of mass/cell.

**Figure 5.**
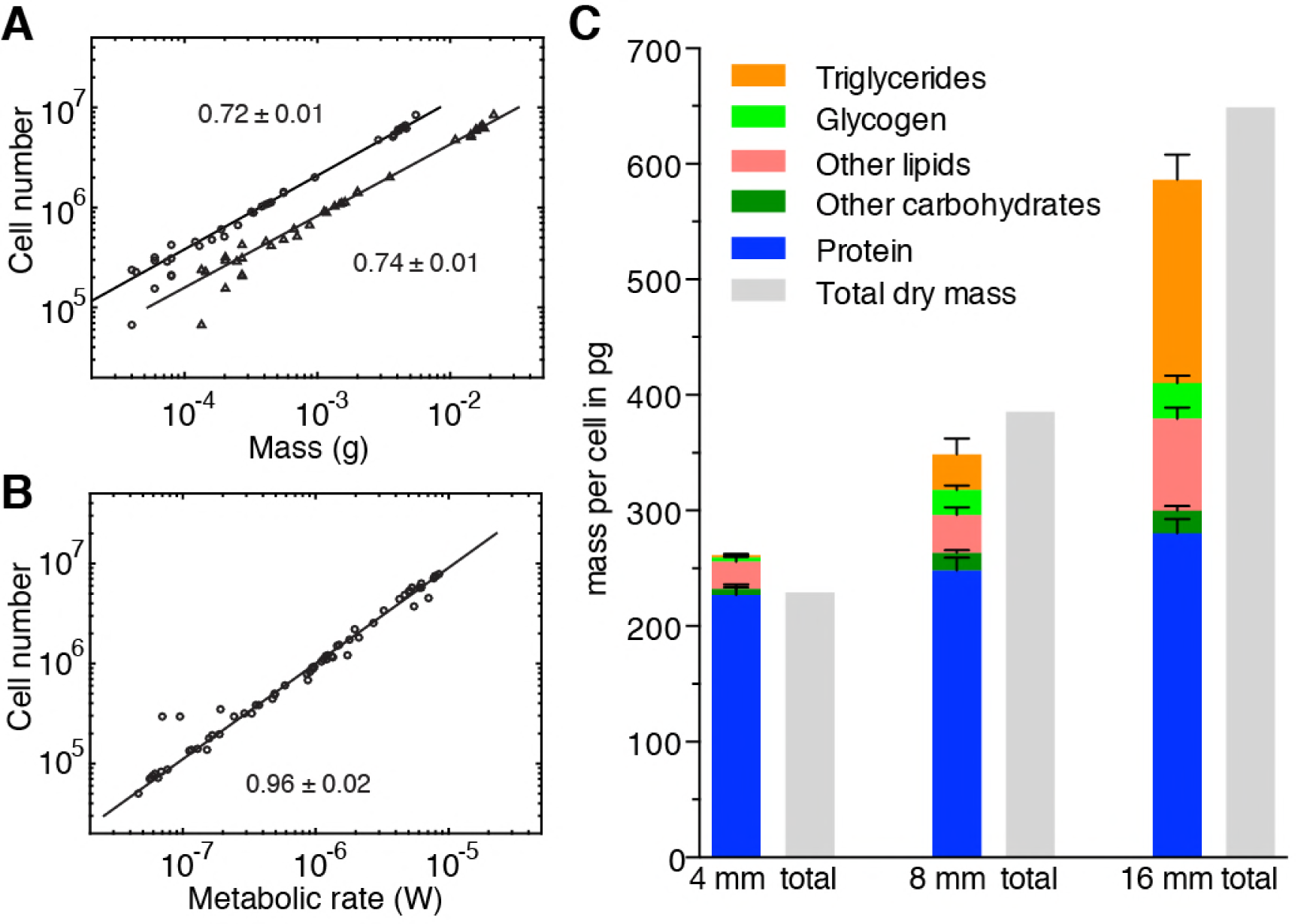
Size-dependent energy storage explains Kleiber’s law scaling. (**A**) Cell number versus dry mass (circles) or wet mass (triangles) based the data from *Figure 3 – figure supplement 3A*. Cell numbers were converted from area using the *N*/*A* scaling law (*Figure 2B*). Dry and wet mass conversion is given by *Figure 1B*. Scaling exponents ± standard errors were derived from respective linear fits and represent the exponent *b* of the power law *y = ax*^*b*^ (**B**) Cell number versus metabolic rate, derived from *Figure 1B* with scaling laws of *Figure 2B* and *Figure 3 – figure supplement 3A*. The scaling exponent ± standard error was derived from respective linear fits and represents the exponent *b* of the power law *y = ax*^*b*^. (**C**) Mass composition (coloured) and total dry mass (grey) per cell in animals of the indicated body length. Triglyceride and glycogen measurements are taken from *Figure 4B* and *4D*. Quantification of other (polar and non-polar) lipids is based on the mass-spectrometry data from *Figure 4B* (n=5 biological replicates; *p*_adj_=0.1720 (no significance) 8 vs. 4 mm, *p*_adj_<0.0001 16 vs. 4 mm, *p*_adj_<0.0001 16 vs. 8 mm; 2 technical replicates). Other carbohydrates represent total carbohydrate minus glycogen. n=4 biological replicates (independent experiments), 40 pooled 4 mm, 20 8 mm, 8 16 mm long animals; *p*_adj_=0.0047 8 vs. 4 mm, *p*_adj_=0.0005 16 vs. 4 mm, *p*_adj_*=*0.2790 16 vs. 8 mm; 3 technical replicates. Protein content was measured colorimetrically. n=4 biological replicates (independent experiments), 44 pooled 4 mm, 10 8 mm, 10 16 mm long animals; *p*_adj_=0.0020 8 vs. 4 mm, *p*_adj_<0.0001 16 vs. 4 mm, *p*_adj_=0.0007 16 vs. 8 mm) (see also *Figure 5 – figure supplement 1*). Significance was assessed by one-way ANOVA followed by Tukey’s post-hoc test. All values were normalised to the total cell number using the previously established length-area (*Figure 1 – figure supplement 1D*) and *N*/*A* (*Figure 2B*) scaling laws. Total dry mass was independently measured (*Figure 3 – figure supplement 3A*) and correlated with length using the length-area relationship (*Figure 1 – figure supplement 1D*). All values are shown as mean ± standard deviation. See *Figure 5 – source data 1* for numerical data and statistics.

To quantitatively assess the contributions of energy stores to the mass/cell scaling exponent and thus to Kleiber’s law, we analysed the composition of the dry mass in small, medium and large animals. In addition to storage lipids and glycogen, we quantified total protein (*Figure 5 – figure supplement 1A*), non-glycogen carbohydrates (*Figure 5 – figure supplement 1B-C*) and other polar and non-polar lipids (*Figure 4 – figure supplement 1B*). In comparison with the 8 and 88-fold increase of glycogen and TG contributions to dry mass/cell, the relative contribution of protein, other polar/non-polar lipids and non-glycogen carbohydrates varied less between small and large animals (*Figure 5C*). Our quantitative assays further allowed us to assess the absolute mass contribution of each compound class to the size-dependent dry mass increase and thus to the origins of the ¾ exponent. Intriguingly, the latter was largely explained by the mass of triglycerides and glycogen, with additional minor contributions from other carbohydrates, polar/non-polar lipids and protein (*Figure 5C*). Overall, our results therefore demonstrate that size-dependent energy storage causes Kleiber’s law scaling in *S. mediterranea.*

## Discussion

Here, we make use of the dramatic and reversible body size changes in the planarian *S. mediterranea* to probe the physiological basis of Kleiber’s law. Contrary to comparative approaches that generally start from *a priori* assumptions and subsequently examine large multi-species data sets for supporting evidence, our finding that planarians obey Kleiber’s law scaling allows us to bring an experimental approach in a model system to bear on the problem. Growth/degrowth in asexual planarians (e.g., the strain examined here) reversibly scales the size of a fully developed adult body plan. This likely exposes size-dependent physiological constraints more clearly than in other animals, where ontogenic growth is often accompanied by developmental changes. Our combination of quantitative and theoretical analysis of growth and degrowth dynamics demonstrate that Kleiber’s law scaling in planarians originates not from a size-dependent decrease in specific metabolic rate, but from a size-dependent increase in average cellular mass. Further, our demonstration that the ¾ exponent is largely caused by size-dependent lipid and glycogen stores firmly ties the physiological origins of planarian Kleiber’s law scaling to energy metabolism. To our knowledge, our results provide a first experimentally founded demonstration of the physiological origins of Kleiber’s law scaling in an animal species.

The fact that we find the cellular metabolic rate to be size-independent contrasts with multiple proposals that envisage the origins of Kleiber’s law in a size-dependent decrease of cellular metabolic activity, e.g., by decreasing mitochondrial density (Smith, 1956). However, our use of microcalorimetry as a pathway-independent read-out of the post-absorptive metabolic rate leaves open the possibility of size-dependencies in the use of specific metabolic pathways. In this context, it is interesting to note that the ratio between our experimentally measured gross energy content (by bomb calorimetry) and net energy usage (modelling of growth dynamics data on basis of paradigm 3; *Figure 3B*) remains constant across the entire size range (*Figure 3F*). This entails that the metabolic processes that assimilate ingested food or catabolise energy stores are largely size-independent (constant food conversion rate of 2.6, in line with other aquatic organisms (Tacon & Metian, 2008); *Figure 3 – figure supplement 3F*). Instead, what changes in a size-dependent manner is the relative proportion of organismal energy resources that is channelled into the formation of metabolically active cells versus metabolically inert energy stores. While small planarians “invest” largely in new cells and little into stores, large animals predominantly store ingested food energy and produce few new cells. Allometric scaling of fat content with mass in mammals (Scale exponent 1.19 (Calder, 1984; Pitts & Bullard, 1968)) raises the possibility that similar physiological trade-offs may contribute to *P*/*M* allometries in other animals.

Our demonstration of size-dependent energy storage as physiological cause of Kleiber’s law in planarians also narrows the quest for a quantitative understanding of the ¾ scale exponent. Interestingly, the trade-off between energy storage and addition of new cells converges on a central premise of the Dynamic Energy Budget (DEB) theory, which can derive the ¾ exponent out of the assumption of surface-limited energy store mobilization (Maino, Kearney, Nisbet, & Kooijman, 2014). However, planarians assimilate and distribute metabolic energy via the branched tubular network of their intestine (Forsthoefel, Park, & Newmark, 2011) (termed “gastrovasculature”). This also makes the intrinsic transport capacity limitations of space-filling fractal networks a possible origin of the ¾ exponent, as per the WEB theory (Goeffrey B West et al., 1997). Further, it is also conceivable that neither process is rate-limiting and that size-dependent energy storage emerges as a consequence of a size-dependence of stem cell division probabilities, for example. Importantly, our results establish an experimental system for the systematic experimental evaluation of these and other theories and thus also the mechanistic basis of Kleiber’s law.

## Acknowledgements

We thank N. Alt, J. Richter, J. Ferria, A. Mishra, M. Toth and I. Smith for size quantifications, K.-O. Linde and IKA Werke GmbH & Co. for microcalorimetry support, Passant Atallah (IPF Dresden) and Michal Surma (Lipotype GmbH) for support with the animal mass measurements and H. Andreas and S. von Kannen for technical support. We thank the CMCB technology platform TU Dresden (EM and Histology) and the following MPI-CBG core facilities for their support: Cell technologies, Technology development studio, and scientific computing.

## Author Contributions

Conceptualization: A.T., S.W., O.F., J.C.R., B.M.F., F.J.; methodology: A.T., O.F., S.W., Y.Q, O.K.; formal analysis: S.W., A.T., O.F., O.K.; investigation: A.T., O.F., S. W., J.P., O.K., Y.Q.; software: S.W.; resources: J.C.R., K.F., A.S., F.S.; data curation: S.W., A.T., O.F., O.K.; writing – original draft: A.T., S. W., O.F., J.C.R.; writing – review & editing: J.C.R., F.J., B.M.F., A.T., S.W., O.F., J.P., K.F., O.K., A.S.; visualization: A.T., S.W., O.F.; funding acquisition: J.C.R., F.J., B.M.F.; K.F. All authors read and approved of the manuscript.

## Data and materials availability

All data on which the conclusions of this paper are based are presented in the figures, figure supplements or source data that was submitted with this manuscript.

## Materials and Methods

### Fitting of power laws

Power law exponents were obtained from linear fits (robust regression using a bisquare weighing function, “robustfit” function in MATLAB) in the log-log plot. We only directly fitted the measured data. If a data set is derived from several measurements (e.g. metabolic rate vs. wet mass is derived from measurements of metabolic rate vs. dry mass and dry mass vs. wet mass), the power law estimate is computed from the original fits of the individual measurements. The respective standard error is obtained via error propagation.

### Animal husbandry

The asexual (CIW4) strain of *S. mediterranea* was kept in plastic containers in 1X Montjuïc salt water (1.6 mM NaCl, 1.0 mM CaCl_2_, 1.0 mM MgSO_4_, 0.1 mM MgCl_2_,0.1 mM KCl, 1.2 mM NaHCO_3_) with 25 mg/L gentamycin sulfate. The animals were fed homogenized organic calf liver paste and were fed at least one week prior to all experiments if not otherwise indicated. Animals were kept at 20 °C before and during experiments.

### Measurement of planarian body size

Movies of gliding planarians were taken with a Nikon Multizoom AZ 100M (0.5x objective) using dark field illumination (facilitates planarian body segmentation). The following camera (DS-Fi1) settings were used: frame rate 3 Hz, exposure time 6 ms, 15 s movie length, 1280 × 960-pixel resolution, conversion factor 44 pixel/mm. Animals were placed one at a time inside a Petri dish and typically 1 - 4 movies taken, depending on the animal’s behaviour. Movies were converted from AVI to MP4 format using Handbreak to reduce the file size. Movies were subsequently analysed using custom-made MATLAB software (MathWorks, Natick, Massachusetts, USA). Typically, those frames were analysed in which the animals were gliding in a straight line (typically 10 frames). See also (Werner et al., 2014).

### Microcalorimetry

Size-matched planarians were placed inside 4 ml glass ampoules (TA Instruments, Cat. No.: 24.20.0401) partially filled with 2 ml of planarian water and supplemented with 10 mM HEPES for improved buffering. No HEPES was used in 22 out of 82 samples, however, no difference in animal health and/or heat generation was observed (data not shown). The ampoules were sealed with aluminium Caps (TA Instruments, Cat. No.: 86.33.0400) using a dedicated crimping tool (TA Instruments, cat. #: 3339). The measurements were performed in a multichannel microcalorimeter (TAMIII, TA Instruments), whereby 12 samples were measured simultaneously including 1-2 controls without animals. The ampoules were first inserted half way and kept in this position for 15 min in order to equilibrate with the temperature inside the device. Then, ampoules were placed completely inside the respective channels whereby they were sitting on top of a thermoelectric detector that measured the heat production in relation to an oil bath, which was kept at a constant temperature of 20 °C. Before the actual measurements, the system was left to equilibrate for another 45 min. The measurements lasted between 2-3 days. Animal behaviour was not controlled and the animals were able to freely move inside the ampoule. Immediately after the metabolic rate measurements, animal dry mass was determined by drying over night at 60 °C either on weighing paper or inside 0.5 ml tubes and subsequent weighing on a microbalance (RADWAG MYA 5.2Y). The mass per animal was obtained by dividing the collective mass by the number of animals.

### Cell counting based on Histone H3 protein quantification

Generating standard curves for converting Histone H3 content into cell number: cells from 15 animals (length 5-8 mm) were dissociated and counted out by FACS essentially as previously described (Tejada-Romero, Evans, & Aboobaker, 2012). Following enzymatic digestion of the tissue, the resulting cell suspension was filtered through a CellTrics 50 µm mesh (Partec, Cat. No.: 04-0042-2317) and incubated in Hoechst (33342) for 1.5 h on a rotator. Subsequently, cells were pelleted once (700 rpm, 10 min) and the supernatant replaced with fresh CMFH. The volume was adjusted to obtain a cell concentration suitable for FACS (typically 1-5·10^6^ cells/ml). Following cell sorting, cells were kept on ice until further processing. Cells were counted with a FACS ARIA III cell sorter (Beckton Dickinson) with standard filter settings and sorted into 2 ml tubes. Typically, 10^5^ cells were sorted per tube. Following FACS, cells were frozen at - 80 ° C until further use.

Determination of total cell number in different-sized planarians using quantitative Western blotting: plan area was measured using above-mentioned method (see also *Figure 2 – figure supplement 1A-C*). Subsequently, individual animals were lysed in 6 M Urea, 2% SDS, 130 mM DTT, 1 µg/ul BSA, 1 µg/ul BSA-AlexaFluor680 conjugate (ThermoFisher Scientific, Ca. No.: A34787), protease inhibitor cocktail and ≥ 2.5 U/ml Benzonase Nuclease (SIGMA, Cat. No.: E1014). Lysis was allowed to proceed for 1 - 1.5 h at room temperature, remaining tissue pieces were completely lysed by tapping the tubes and vortexing. Meanwhile, the cells for the standard curve (see above) were lysed by directly applying the lysis solution onto the frozen cells. Protein concentrations were measured in 1:5 or 1:10 dilutions using a NanoDrop spectrophotometer (Thermo Fisher Scientific) (absorbance at 280 nm). Finally, the samples were mixed with 4x Laemmli buffer (4x stock: 400 mM DTT, 200 mM Tris-HCl, 8% SDS, 40% glycerol, 0.5 mg/ml Bromophenol Blue) and incubated for 10 min at 60 °C before spinning down at 13000 rpm for 5 min. The samples were run on NuPAGE Novex 4-12% Bis-Tris protein gels (Invitrogen, Cat. No.: NP0322BOX) in 1x MOPS running buffer (ThermoFisher Scientific, Cat. No.: NP0001). The loaded volumes for the standard curve corresponded to 15000, 22500, 30000, 37500 and 45000 cells (linear signal range) and the volume of the whole-animal lysates was corresponding to 50 µg of protein, ensuring that the samples were lying within the range of the standard curve. 4 technical replicates were carried out per experiment (analysis of 5 individual animals) by running 2 chambers with 2 gels each at 140 mA for 1 h. Proteins were transferred onto Whatman Protran nitrocellulose membrane (SIGMA, Cat. No.: Z613630) for 2 h in transfer buffer (20% MeOH/1x MOPS). Membranes were blocked for 1 h at room temperature and continuous agitation in 1x TBS-T (10 mM Tris base, 150 mM NaCl, 0.1% (w/v) Tween-20, pH 7.4) and 5% (w/v) nonfat dry milk. Afterwards, membranes were incubated over night at 4 °C with anti-Histone H3 antibody (Abcam, Cat. No.: ab1791) followed by at least 3 washes in TBS-T for 10 min. Membranes were then incubated with a fluorophore-conjugated secondary antibody (anti-rabbit IRDye 680LT, LICOR, Cat. No.: 926-68023) diluted 1:20000 in blocking solution followed by extensive washing in TBS-T (1x 5 min, 3x 10 min) and one final wash step in TBS (10 min). Afterwards, membranes were dried at room temperature for at least 1 h and imaged on an Odyssey SA Li-Cor Infrared Imaging System (LICOR). The relative fluorescent band intensity was quantified using the gel-analysing tool in Fiji (Schindelin et al., 2012). The fraction of cells from whole-animal lysates loaded onto the gel was calculated from the standard curve on each blot separately. The total number of cells in the animals was calculated as follows: number of cells loaded/volume loaded x total volume of original lysate. The obtained values were finally averaged over all 4 technical replicates.

### Image-based cell counting

First, plan area of individual animals was measured using above-mentioned method (see also *Figure 2 – figure supplement 1A-C*). For cell dissociation, individual animals were placed inside maceration solution (Romero & Baguñà, 1991) (acetic acid, glycerol, dH_2_O at a ratio of 1:1:13 including 1 µg/ml BSA + 10 µg/ml Hoechst 33342, no methanol) and the total volume adjusted according to animal size. The solution also contained typically about 1.3 × 10^6^ fluorescent beads/ml (FluoSpheres Sulfate Microspheres, 4 µm, red fluorescent 580/605 nm, ThermoFisher Scientific, Cat. No.: F8858) the concentration of which was determined with a Neubauer chamber for each experiment (including 10-18 animals). Dissociation was allowed to proceed at room temperature for about 15 min after which cells of remaining tissue clumps were further dissociated by taping and vortexing. Per animal, 2 µl drops of the cell suspension were pipetted into 6-10 wells of a glass bottom 96-well plate (Greiner, Cat. No.: 655090) and the drops dried over night at room temperature. Subsequently, the entire drops were imaged on an Operetta high-content imaging system (PerkinElmer). The number of cells and beads were automatically counted using an imaging pipeline built in CellProfiler (Carpenter et al., 2006) (*Figure 2 – figure supplement 2*). The total number of cells was calculated from each separate well/drop by the following formula: sum of cells in analysed images/sum of beads in analysed images x known total number of beads in original cell suspension. For each animal, the calculated total cell number was averaged across 9-10 wells.

### Measurement of energy content using a bomb calorimeter

Size-matched planarians were placed inside a combustion crucible and lyophilized overnight in a lyophiliser (Heto LyoLab 3000). Then, the samples were weighed on an analytical balance (Sartorius Entris, readability: 0.1 mg) and the mass per animal was obtained by dividing the collective mass by the number of animals – thus, allowing further conversion into organismal cell numbers. Afterwards, the combustion enthalpy was measured by combustion in the presence of high pressure O_2_ inside a bomb calorimeter (IKA C 6000 global standards) running in adiabatic mode. Benzoic acid pellets (IKA C723, Cat. No.: 0003243000) were used as a standard for calibration as well as a burning aid for the samples. In between lyophilizing and combustion, the samples were kept inside a drying chamber to prevent humidification.

### Dry and wet mass measurements

To obtain the dry mass versus area and dry mass versus length scaling laws, the plan area of individuals animals was measured using aforementioned method. Afterwards, animals were individually placed on round pre-weighed glass cover slips and dried over night at approximately 60 °C. Subsequently, each animal was weighed 3 times on an analytical microbalance (Sartorius Research 210 P) to obtain an average mass value. Wet mass was measured by removing as much of residual water as possible while individual animals were placed inside a 0.5 ml tube. After further exposing the animals to air for 30-40 min to evaporate remaining water outside of the animal, animals were weighed on a microbalance (RADWAG MYA 5.2Y).

### Food intake assay

Plan area of individual animals (two and three weeks starved) was measured using the above-mentioned method (see also *Figure 2 – figure supplement 1A-C*). Planarians were fed with organic homogenized calf liver paste, which was mixed with about 6.5*10^5^ per 1 µl liver red fluorescence beads (FluoSpheres Sulfate Microspheres, 4 µm, fluorescent 580/605 nm, ThermoFisher Scientific, Cat. No.: F8858) coated in 1mg/ml BSA. Single animals (or for calibration 2 µl of liver/beads mix) were dissociated into single cells in maceration solution (see above) containing 0.1% Tween-20 and about 300/µl yellow-green fluorescence beads (FluoSpheres Sulfate Microspheres, 4 µm, fluorescent 505/515 nm, ThermoFisher Scientific, Cat. No.: F8859) for volume normalization (see further below). 1 µl drops of the animal and liver macerates as well as from maceration solution only were distributed into 10 wells of a glass bottom 96-well plate (Greiner, Cat. No.: 655090) and dried over night at room temperature in the dark. Whole drops were imaged on an Operetta high content imaging system (PerkinElmer) and the number of red and yellow-green beads were automatically counted using CellProfiler (Carpenter et al., 2006). The volume of liver eaten per animal was calculated as follow:

1. Total number of red beads per 1 animal = Number of red beads in 1 µl drop of worm suspension x Total volume of original maceration solution
2. Total number of red beads per 1 µl liver = (Number of red beads in 1 µl drop of liver suspension **/** 2) x Volume of maceration solution
3. Volume of liver eaten per animal = Total number of red beads per 1 animal **/** Total number of red beads per 1 µl liver

To account for possible pipetting errors leading to variation in drop volumes, the volume of liver eaten per animal was normalized to the ratio between yellow-green beads in the drops of the animal macerate and in the drops of maceration solution only.

### Lipid droplet stain

Two weeks starved small worms were killed in 5 % N-Acetyl-Cystein (NAC) and large worms in 7.5 % NAC (5 min at room temperature) and fixed in 4 % PFA for 2 days at 4°C. Fixed worms were embedded in 4 % low-melting-point agarose and sectioned using a vibratome (100 µm, Leica, Germany). Sections were treated with 0.5 % Triton X-100 in PBS for 2h and incubated in PBS with lipid droplet dye LD540 (kind gift from Christoph Thiele, Bonn) (0.5 µg/ml) and DAPI (1 µg/ml) overnight at room temperature. After thoroughly washing with 0.3% Triton X-100 in PBS and a short rinse in PBS, the sections were optically cleared with the slightly modified SeeDB protocol (Ke, Fujimoto, & Imai, 2013) as follows: Sections were incubated sequentially with increasing concentrations of aqueous fructose solution (25 % for 4 h, 50 % for 4 h, 75 % and 100 % fructose for overnight) and finally with the saturated fructose solution overnight. All steps were carried out at room temperature. The sections were mounted on glass slides with the SeeDB solution and confocal images were taken on a Zeiss LSM 700 inverted microscope (20x objective, Zeiss Plan-Apochromat, 0.8 numerical aperture) using 80 % 2,2’-Thiodiethanol (Staudt, Lang, Medda, Engelhardt, & Hell, 2007) as immersion media.

### Lipid extraction and quantification by shotgun mass spectrometry

Synthetic lipid standards were purchased from Avanti Polar Lipids, Inc. (Alabaster, AL, USA). Stocks of internal standards were stored in glass ampoules at −20°C until used for lipid extraction. Planarians of different size (40 small, length ~ 4 mm; 20 medium, ~ 8 mm; and 6 large, ~ 16 mm) were pooled and homogenized in ice-cold isopropanol mixed with acetonitrile (1:1). Analysis of lipid extracts obtained under different homogenization conditions to prevent TG degradation (*Figure 4 – figure supplement 1*) was performed on HPTLC silica gel plates (Merck, Cat.No.: 105633) with the solvent system n-hexane/diethylether/acetic acid (70:30:1, vol/vol/vol). Lipids were visualized by spraying plates with 3 g cupric acetate in 100 ml of aqueous 10 % phosphoric acid solution and heating at 180 °C for 10 min. Protein amount in the homogenates was determined by BCA. 50 µg of total protein was extracted with MTBE/MeOH as described in (Sales et al., 2016; Sales, Knittelfelder, & Shevchenko, 2017; Schuhmann et al., 2012). Briefly, 700 µl of 10:3 MTBE/MeOH containing one internal standard for each lipid class was added to the dried homogenates. Samples were vortexed for 1h at 4 °C. Phase separation was induced by adding 140 µl of water and vortexing for 15 min at 4 °C, followed by centrifugation at 13400 rpm for 15 min. The upper phase was collected, evaporated and reconstituted in 600 µl of 2:1 MeOH/CHCl_3_. 15 µl of total lipid extract was diluted with 85 µl 4:2:1 IPA/MeOH/CHCl_3_ containing 7.5 mM ammonium formate for mass spectrometric analysis. For the measurement of phosphatidylserines (PS), 15 µl of lipid extract were diluted with 85 ul 4:1 EtOH/CHCl_3_ containing 0.1% triethylamine.

Mass spectrometric analysis was performed on a Q Exactive instrument (Thermo Fischer Scientific, Bremen, Germany) equipped with a robotic nanoflow ion source TriVersa NanoMate (Advion BioSciences, Ithaca, NY, USA) using nanoelectrospray chips with a diameter of 4.1 µm. The ion source was controlled by the Chipso?t 8.3.1 so?tware (Advion BioSciences). Ionization voltage was + 0.96 kV in positive and - 0.96 kV in negative mode; backpressure was set at 1.25 psi in both modes by polarity switching (Schuhmann et al., 2012). The temperature of the ion transfer capillary was 200 °C; S-lens RF level was set to 50 %. Each sample was analysed for 5.7 min. FTMS spectra were acquired within the range of m/z 400–1000 from 0 min to 1.5 min in positive and within the range of m/z 350–1000 from 4.2 min to 5.7 min in negative mode at a mass resolution of R m/z 200 = 140000, automated gain control (AGC) of 3 × 10^6^ and with a maximal injection time of 3000 ms. Free cholesterol was quantified by parallel reaction monitoring FT MS/MS within runtime 1.51 to 4.0 min. For FT MS/MS micro scans were set to 1, isolation window to 0.8 Da, normalized collision energy to 12.5%, AGC to 5 × 10^4^ and maximum injection time to 3000 ms. PS was measured for 1.5 min in an additional acquisition in negative FTMS mode with optimized nanoMate parameters (backpressure 1.00 psi and voltage – 2.00 kV). All acquired data was filtered by PeakStrainer (https://git.mpi-cbg.de/labShevchenko/PeakStrainer/wikis/home) (Schuhmann et al., 2017). Lipids were identified by LipidXplorer so?tware (Herzog et al., 2012). Molecular Fragmentation Query Language (MFQL) queries were compiled for PC, PC O-, LPC, LPC O-, PE, PE O-, LPE, PI, LPI, PA, LPA, PS, SM, TG, DG, Cer, Chol, CE lipid classes (see Table 1 for meaning of all abbreviations). The identification relied on accurately determined intact lipid masses (mass accuracy better than 5 ppm). Lipids were quantified by comparing the isotopically corrected abundances of their molecular ions with the abundances of internal standards of the same lipid class. The amount of lipids per animal was calculated based on the known volume of homogenization buffer and the known number of animals. Lipid amounts were normalized to cell number using the previously established scaling relationship between cell number and area (*Figure 2B*) and between length and area (*Figure 2 – figure supplement 1D*).

### Histological staining for glycogen on planarian cross sections

Fixation: two weeks-starved small (~ 4mm) and large (13mm −16mm) animals were anesthetized and relaxed for 5 min on ice by supplementing chilled planarian water with 0.0485% w/v Linalool (Sigma, L2602). Planarians were fixed in cold alcoholic Bouins fixative (15ml Picric acid (saturated alcoholic solution, TCS Biosciences, Cat. No.: HS660), 12 ml 32 % PFA, 2 ml glacial Acetic acid and 15 ml Ethanol) for overnight at 4°C and washed with 70 % Ethanol for following two days.

Paraffin embedding and sectioning: Fixed animals were dehydrated by alcohol-xylene series (1x 10 min in 70 % ethanol and 2x for 30 min in 96 %, 100 % ethanol and xylene, respectively). Xylene was replaced by melted paraffin at 60 °C, which was exchanged three times, after 30 min, after several hours overnight and again after 30 min, which was followed by embedding. Cross-sections of 10 µm thickness were obtained using a microtome (Thermofisher Scientific, Microm HM355S). The sections were dewaxed and hydrated by xylene-ethanol series (2x 10 min Xylene, 2x 1min 100 %, 96 % and 1×1 min 70 %, 40 %, ethanol and water). Prior to staining, one of two adjacent sections was treated (for 2 h, at 37 °C) with 0.2 N acetate buffer (pH 4.8) containing amyloglucosidase (0.03 U/µl) (Sigma A1602), while the other section with buffer only. By rinsing the sections with water, the digested glycogen was washed out on the section treated with amyloglucosidase but not on the section without enzyme treatment.

For glycogen visualization, we used Best’s Carmine staining method. The Carmine stock and - working solutions (Carmine (C.I. 75470) Carl Roth, 6859.1) as well the differentiating solution were prepared as described in Romeis - Mikroskopische Technik (Mulisch & Welsch, 2010). The sections were treated for 10 min with Carmine working solutions following by differentiating solution 2x for 1 min. Sections were briefly rinsed with 80 % ethanol and treated 2x for 1 min with 100 % Ethanol and 2x for 2 min with Xylene and mounted in Cytoseal(tm)XYL (Richard-Allan Scientific; 8312-4). Stained sections were imaged with an Olympus BX61 Upright Microscope with 5x and 20x objectives.

### Glycogen assay

Two weeks-starved animals were homogenized in water (40 worms of 4 mm length in 0.5 ml, 20 worms of 8 mm in 1ml and 10 worms of 16 mm in 1ml) using zirconia/silica beads (1.0 mm diameter, Carl Roth GmbH+Co.KG, Cat.No:11079110z) at 4°C for 10 min. After brief centrifugation, the samples were flash frozen in liquid nitrogen and sonicated (Covaris S2 Sonicator) for 1 min. The homogenate was used for glycogen and total carbohydrate quantifications. The glycogen quantification method was adapted to planarians based on a protocol for *Drosophila* larvae from the C. Thummel lab (University of Utah). Heat-treated homogenate (70°C, 10 min) was centrifuged at 13400 rpm for 2 min and the supernatant was taken for the measurements. The extracted glycogen was digested to glucose by amyloglucosidase treatment (Sigma, Cat. No.: A1602) (0.015 U/µl of 0.2 M acetate buffer, pH 4.8) for 2 h at 37 °C. The glucose content was measured using the glucose assay kit (Sigma, Cat. No.: GAGO-20). The assay was performed in black 96-well glass bottom plates (Greiner Bio-One, Cat. No.: 655090) and the absorption spectra was measured using Envision Microplate Reader (Perkin Elmer). Additionally, to assess background levels of free glucose, the supernatant without amyloglucosidase treatment was measured. Planarians do not contain free Glucose at detectable levels (data not shown). Glucose and Glycogen amounts were determined using a standard curve built on a glucose and glycogen dilution series, respectively. Glycogen extraction using hot 30 % KOH (*Figure 4 – figure supplement 1D*) was performed as previously published (Rasouli, Shokri-Afra, & Ostovar-Ravari, 2015).

### Total carbohydrate measurement

Determination of total carbohydrate was carried out on whole homogenates (same as used in glycogen assay) using the phenol-sulfuric acid method. In brief, the homogenate was heated with the 96% H_2_SO_4_ at 90°C for 15 min, mixed with phenol (saturated with 0.1M citrate buffer, pH 4.3, Sigma, Cat. No.: P4682) (Homogenate: H_2_SO_4_: phenol at a ratio of 1:5:5) and distributed into a 96-well plate (Thermo Scientific Nunc, Cat. No: 167008). The absorbance was measured at 492 nm Envision Microplate Reader (Perkin Elmer). Carbohydrate amounts were determined using the glycogen standard curve (see previous section). The amount of glycogen and total carbohydrates per animal was calculated based on the known volume of homogenisation buffer and the known number of animals. Glycogen and carbohydrate amounts were normalised to organismal cell number using the previously established scaling relationship between cell number and area (*Figure 2B*) and between length and area (*Figure 2 – figure supplements 1D*). The non-glycogen carbohydrate amount was calculated by subtracting the determined glycogen from the carbohydrate amount.

### Protein measurements

Planarians of approximately 4, 8 and 16 mm length were chosen and protein amounts were determined using the Pierce 660nm Protein Assay Reagent (ThermoFisher Scientific, Cat. No.: 22660) according to the manufacturer’s instructions. To ensure compatibility with the used lysis solution (see below), the Pierce 660nm Protein Assay Reagent was complemented with Ionic Detergent Compatibility Reagent (ThermoFisher Scientific, Cat. No.: 22663). Planarian lysates were prepared as follows: 44 small (length 4 mm), 10 medium (8 mm) and large (16 mm) animals were placed inside 1.5 ml tubes and rinsed once with dH_2_O. A lysis solution containing 10 M Urea, 2 % s odium dodecyl sulfate (SDS), 130 mM dithiothreitol (DTT), 2.5 µg/ml Benzonase (home-made) and a protease inhibitor cocktail was added and the animals incubated for 10 min followed by homogenisation using a motorized plastic pestle. Volumes of lysis buffer used were 235 µl for small, 335 µl for medium and 2 ml for large animals. Subsequently, lysates were cleared by centrifugation at 13000 rpm for 1 minute. The assay was performed in black 96-well glass bottom plates (Greiner Bio-One, Cat. No.: 655090) and the resulting absorption spectra measured using a FLUOstar Omega Microplate Reader (BMG LABTECH).

### Whole mount in situ hybridization

Whole mount *in situ* hybridization (WISH) was essentially performed as previously described (King & Newmark, 2013; Pearson et al., 2009).

### Statistics

All statistical analyses were carried out using GraphPad Prism version 7.0c for Mac OSX (GraphPad Software, La Jolla, California, USA).

### Software

Excel for Mac (Microsoft, Redmond, Washington, USA) and KNIME(Berthold et al., 2007) (KNIME AG, Zurich, Switzerland) were used for data handling and calculations; GraphPad Prism v7.0c (GraphPad Software, La Jolla, USA) was used for statistical analyses and data visualization; MATLAB (MathWorks, Natick, Massachusetts, USA) was used for planarian body size measurements, theoretical analysis of models, data handling and visualisation; CellProfiler (Carpenter et al., 2006) was used for image analysis; Fiji (Schindelin et al., 2012) was used for Western blot quantification and image processing; Adobe Photoshop CS5 and Illustrator CS5 (Adobe Systems, San Jose, California, USA) were used for image processing and generating figures; the manuscript was prepared for submission using Word for Mac (Microsoft, Redmond, Washington, USA).

## Figure supplements

**Figure 1 – figure supplement 1.**
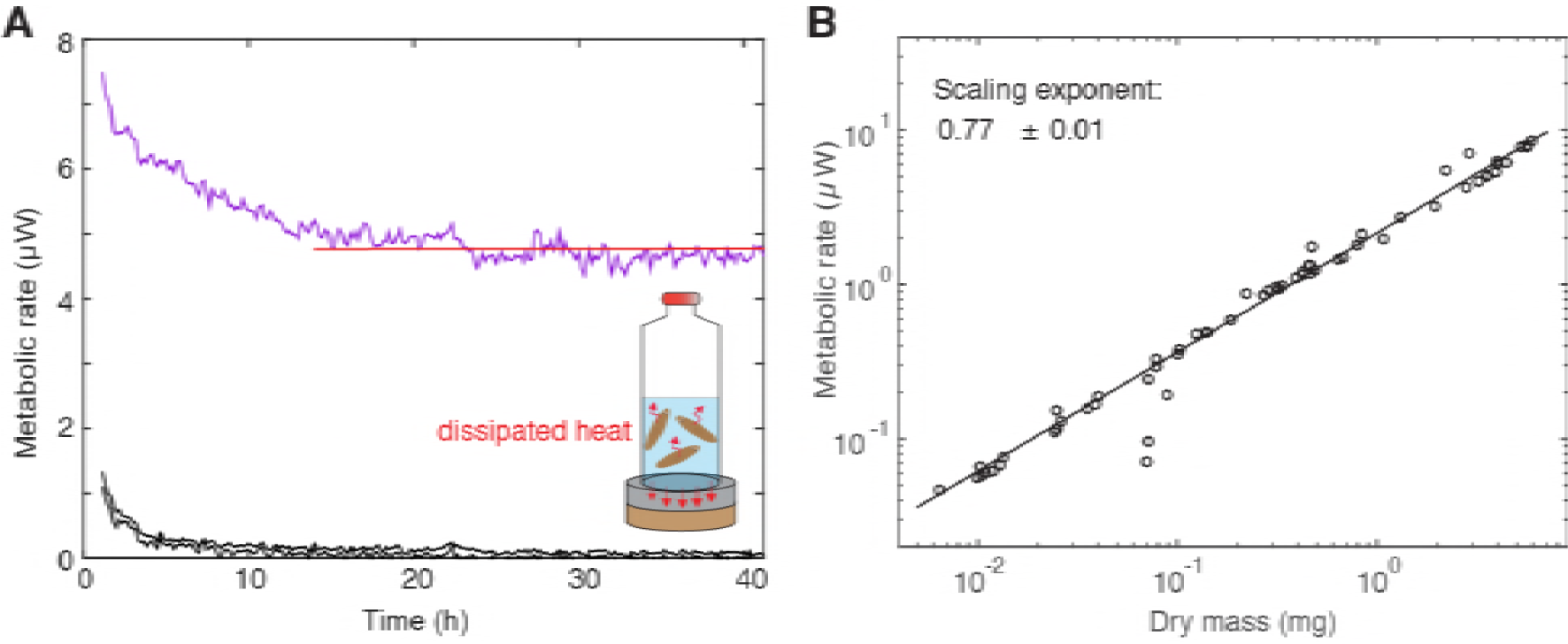
Measurement of metabolic rate. (**A**) Cartoon of a vial with enclosed animals placed on a thermo-electric detector and representative metabolic rate (heat flow) trace with an initial equilibration followed by a stable phase. The thermo-electric detector measures the heat production relative to an oil bath kept at constant temperature. The average metabolic rate was determined by manually fitting a horizontal line (red). The first 10 h were always excluded because of the initial relaxation of the control signal (medium only) which is shown in black. For individual samples, the relaxation time has been estimated to be larger than 10 h. For 21 out of 83 samples, no fit could be obtained due to high fluctuations of the signal. (**B**) Metabolic rate versus dry mass scaling used to plot *Figure 1C*. The scaling exponent ± standard error was derived from a linear fit and represents the exponent *b* of the power law *y = ax*^*b*^. See *Figure 1 – figure supplement 1 – source data 1* for numerical data.

**Figure 2 – figure supplement 1.**
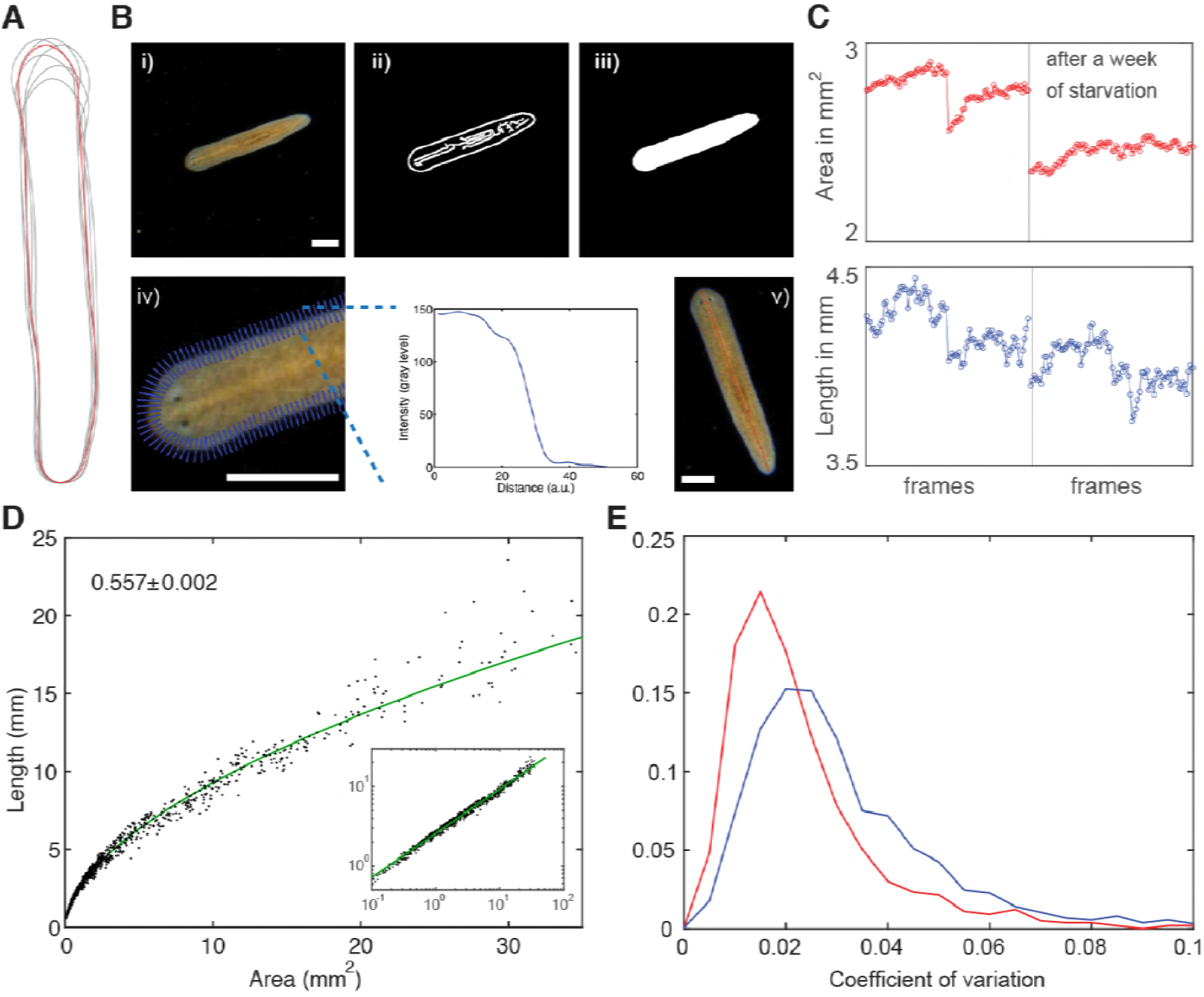
Measurement of planarian body size. (**A**) Variable body outline of one individual extracted from a series of movie frames. (**B**) Pipeline for extracting plan area and length of planarians from movies, see also (Werner et al., 2014). i) raw movie frame, ii) use of Canny filter on background-corrected frames to identify edges, iii) dilation-erosion cycle to fill holes, iv) refinement of animal perimeter by finding steepest intensity across body edge, v) final boundary outline (blue) and midline (red) used to calculate area and length. Scale bars, 1 mm. See *Figure 2 – source data 4* for MATLAB script. (**C**) Area (top) and length traces (bottom) of the same individual separated by one week of starvation (compare left and right box). A drop of plan area and length reflects the expected decrease in body size and shows sensitivity, in particular of the less variable area, to small size changes. (**D**) Length versus area scaling including almost 900 measurements suggests tight regulation of body shape during growth and degrowth and confirms the accuracy of our size quantification method. The scaling exponent ± standard error was derived from a linear fit and represents the exponent *b* of the power law *y = ax*^*b*^. Inset shows the same data plotted on logarithmic axes. (**E**) Histogram of the coefficients of variation (ratio standard deviation/mean) revealing less variability of the plan area compared to body length. See *Figure 2 – figure supplement 1 – source data 1* for data of (D) and (E).

**Figure 2 – figure supplement 2.**
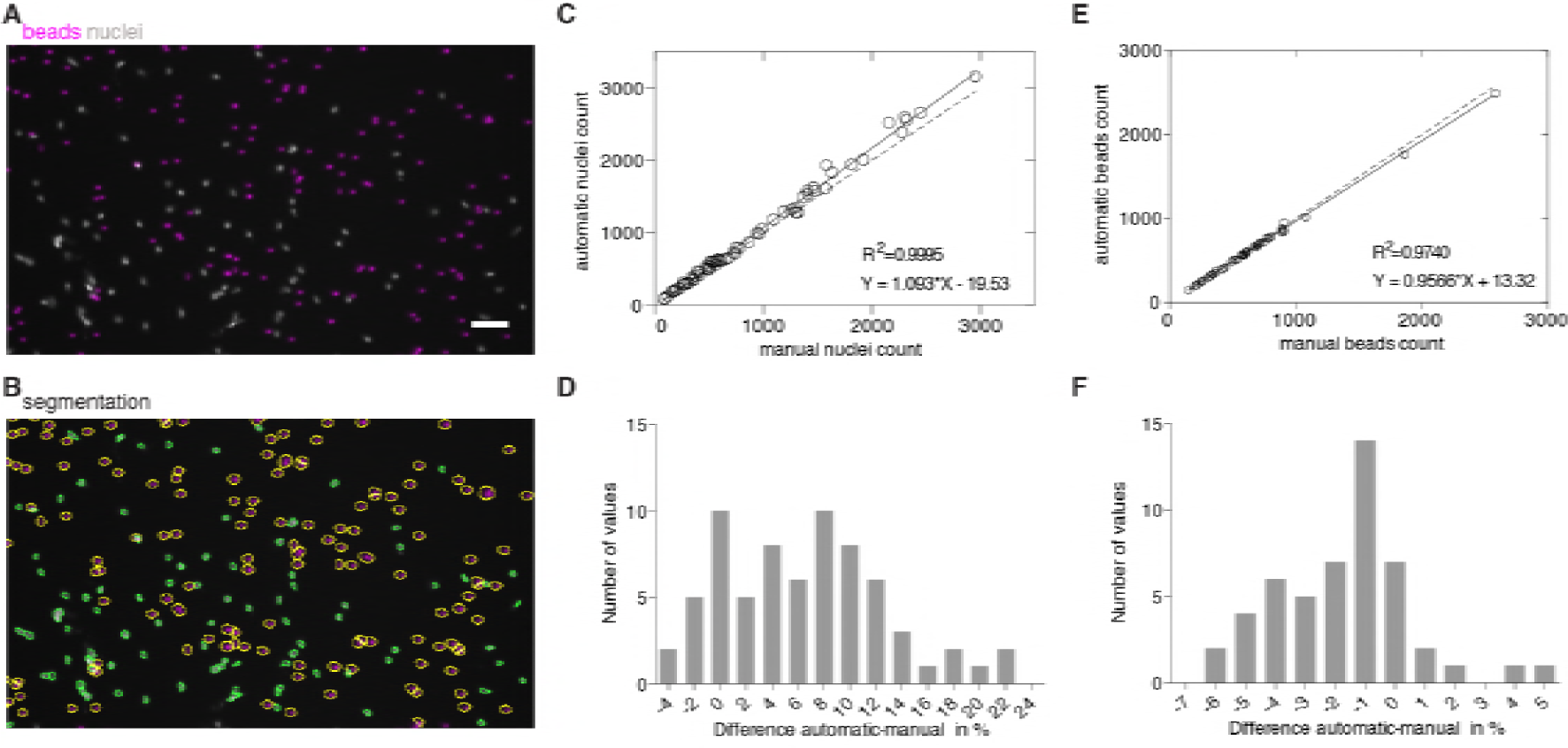
Validation of image-based quantification of organismal cell number. (**A**) Nuclei (grey) and fluorescent beads (magenta) inside a drop. (**B**) Automatic segmentation of nuclei (green contours) and beads (yellow contours). Scale bar, 25 µm. (**C**) The high correlation between automatic and manual counting methods confirms accuracy of nuclear counts (analysis of 69 images from 3 experiments). Solid line, linear regression; dotted line, hypothetical perfect match between automatic and manual counts (*y=x*). (**D**) Histogram showing the (binned) number of measured differences per image between automatic and manual counting (**E**), The high correlation between automatic and manual counting methods confirms accuracy of bead counts (analysis of 50 images from 3 experiments). Solid line, linear regression; dotted line, hypothetical perfect match between automatic and manual counts (*y=x*). (**F)** Histogram showing the number of (binned) measured differences per image between automatic and manual counting. See *Figure 2 – figure supplement 2 – source data 2* for raw images and numerical data of (C)- (F).

**Figure 2 – figure supplement 2.**
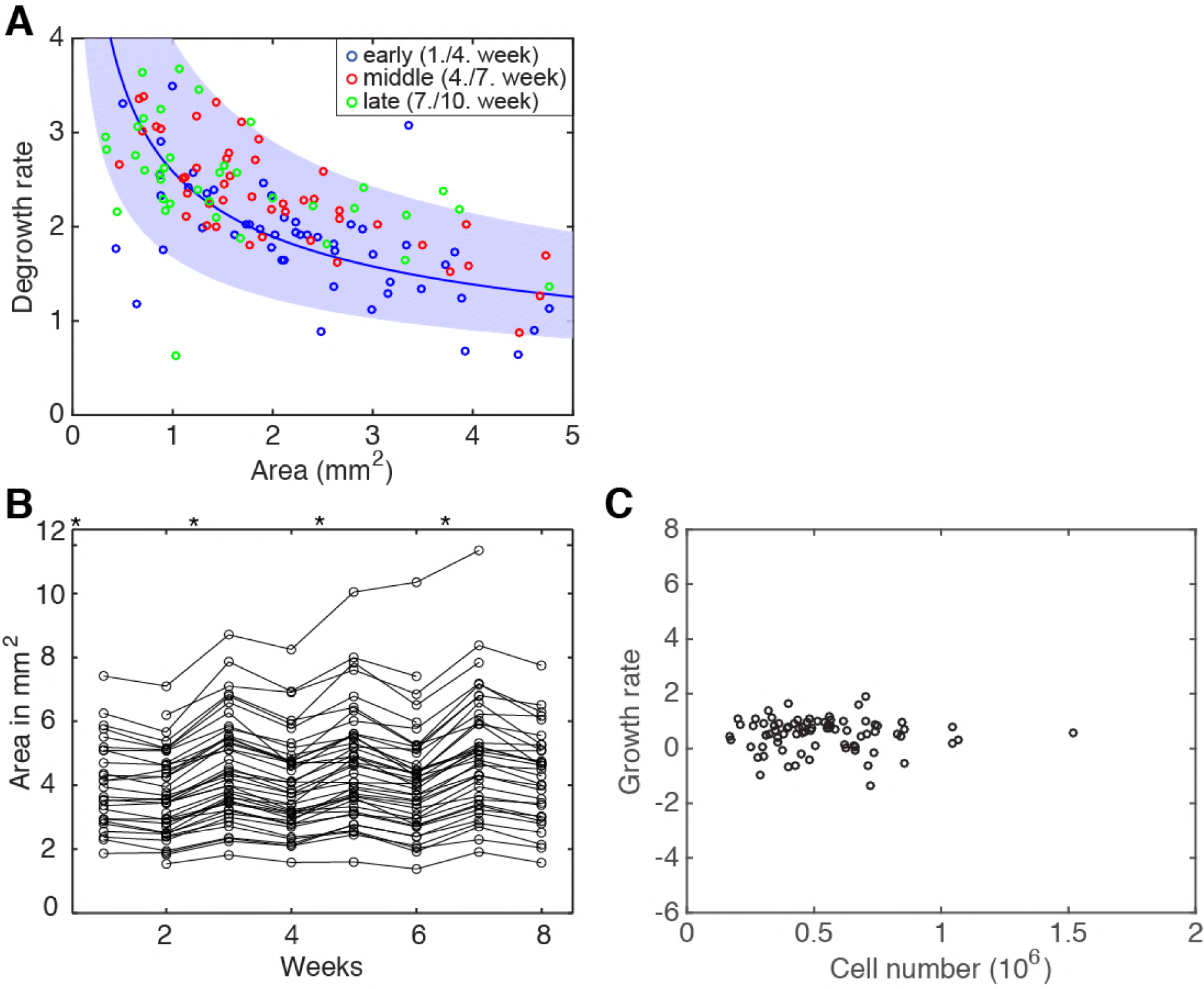
Degrowth rates are independent of feeding history. (**A**) Degrowth rates from Fig. 2E colour-coded according to time since start of food deprival; early (blue), 1-4 weeks; middle (red), 4-7 weeks; late (green), 7-10 weeks. Solid line represents a power law (*y = ax*^*b*^) fit to the early (blue) time points. Blue band represents the 95 % confidence interval for the early (blue) degrowth rates; note that middle (red) and late (green) degrowth rates lie mostly within this interval, hence, are not significantly different from the earliest points (no feeding history). (**B**) Area change of individual animals fed every second week (* indicate feeding time points). Every feeding event results in a small growth peak. While feeding every second week maintains on average a stable body size, an increased feeding frequency would cause addition of small growth peaks resulting in long-term growth. See *Figure 2 – source data 5* for respective data. (**C**) Calculating de-/growth rates based on (B) reveals that feeding every second week, on average, results neither in growth nor in degrowth. Individual data points were calculated by exponential fits to traces in (B) across <u>2</u> time windows and using the *N*/*A* scaling law (*Figure 2B*) to express rates as % change in cell number/day.

**Figure 3 – figure supplement 1 (text only). Implementation of the theoretical model**

The model describes the dynamic changes of total physiological energy *E*, defined as the fraction of energy in the body that can be metabolized and released as heat, see *Figure 3A*. The physiological energy *E* thus decreases due to metabolic heat production *P* and increases due to feeding, where *J* captures the net influx of physiological energy (taking into account a potentially elevated metabolism during feeding): 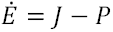. The dot denotes the time derivative. The average energy per cell is computed by dividing the total physiological energy by the total number of cells *N*: *e* = *E*/*N*. Thus, the energy per cell changes according to: 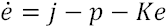, where we define *j* = *J*/*N, p* = *P*/*N* and the growth rate *K* = *Ṅ*/*N*. An increasing cell number decreases the energy per cell.

Paradigm 1 assumes that cell division and cell death directly depend on the energy available per cell *e*. For simplicity, we consider a linear relationship between the growth rate and the energy per cell: *K* = *K*_0_(*e*/*e*_*s*_ − 1), with *K*_+_ being a characteristic rate of growth and degrowth and *e*_*s*_ being the critical energy per cell at which planarians switch between growth and degrowth (*Figure 3 – figure supplement 2A*, dashed line). Thus, we can describe the energy dynamics by *ė* = *j* – *p* – *K*_0_(*e*/*e*_*s*_ − 1)*e*. During starvation, *j* = 0 and the growth rate is decreasing, which requires 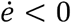 (red curve). The maximum of *ė* is at *e* = *e*_*s*_/2 and from 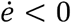 follows that *p* > *e*_*s*_*K*_0_/4. During feeding, where *j* > *p* the curve is shifted upwards and *e* ends up in a growth regime (blue curve). For a constant energy influx *j*, the equation for 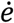 has a stable fixed point 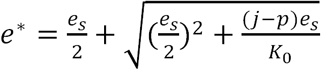 with *e*^*^ > *e*_*s*_ for *j* > 0. Thus, the animal would grow at a constant rate *K*^*^ = *K*_0_(*e*^*^/*e*_*s*_ − 1). In order for the growth rate to decrease with animal size (*Figure 2E*), the energy influx per cell *j*(*N*) must not be constant but has to be a decreasing function of *N*, hence, we choose *j*(*N*) = *j*_0_/(1 + *N*).

*Figure 3 – figure supplement 2B* shows a time course of the organismal cell number *N* when going through several rounds of feeding (blue) and starvation (red), always switching at a certain size, specifically at *N* = 0.5 · 10^6^ cells and *N* = 4.5 · 10^6^ cells (lower and upper dashed lines, respectively). In the beginning of the starvation interval, we see an overshoot where the animal still grows although feeding has stopped. As a result, we observe rather generic growth and degrowth kinetics, independent of initial values for energy and cell number or the feeding scheme, see *Figure 3 – figure supplement 2C*. Any perturbation decays quickly and there is no strong dependence on feeding history.

Paradigm 2 and 3 assume a constant relationship between cell number and physiological energy content of the worm: *E*~*N* and *E*~*N*^*c*^, respectively. In consequence 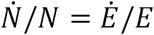 and *Ṅ*/*N* = *Ė*/(*Ec*), respectively, which can be related to metabolic rate and feeding influx via 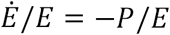 during degrowth and via 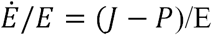 during growth. In paradigm 2, *E*/*N* is constant, therefore both *P*/*N* and *J*/*N* have to depend on *N* to explain the growth and degrowth rates. In paradigm 3, both *P*/*N* and *J*/*N* can be chosen to be constant.

To fit the growth dynamics in *Figure 2C* by paradigm 1, we use the following parameters:

*j*_0_/*e*_*s*_ = 14%/*d, p*/*e*_*s*_ = 1.3%/*d, K*_0_ = 4.3%/*d*, initial conditions *N*(0) = 3 · 10^6^ and *e*(0)/*e*_*s*_ = 1 as well as a switch between feeding and starvation regimes at *N* = 0.05 · 10^6^ and *N* = 6.5 · 10^6^. Yet, several combinations of parameter values can fit the measurement equally well. From a fit of paradigm 2 in *Figure 2C*, we obtain *P*/*E* = 243 *N*^−1.35^%/*d* and *J*/*E* = 214 *N*^−0.28^%/*d*. Finally, from a fit of paradigm 3 to the data, we obtain *E*/*P* = 0.34 *N*^0.38^*d* and *J*/*P* = 3.0, see *Figure 2C*.

**Figure 3 – figure supplement 2.**
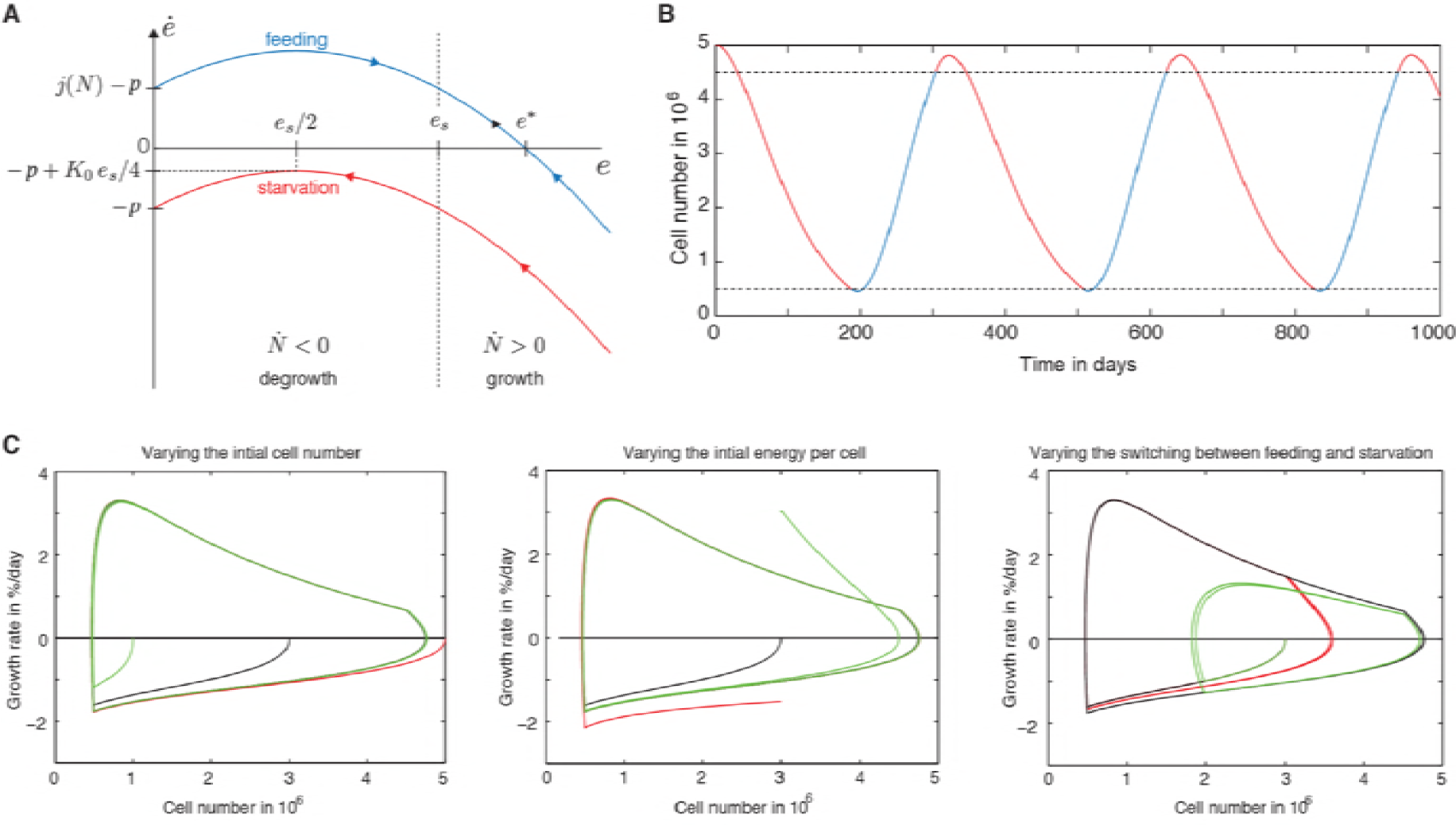
Further explanation of model paradigm 1. (**A**) Dynamic behaviour of the energy content per cell during feeding and starvation in paradigm 1. Graph shows the change of the energy content per cell *e* as a function of the energy content per cell *e* during starvation and feeding. See *Figure 3 – figure supplement 1* for a detailed description of paradigm 1. (**B**) Time course of the organismal cell number *N* when going through several rounds of feeding (blue) and starvation (red), always switching at a certain size, specifically at *N* = 0.5 · 10^6^ cells and *N* = 4.5 · 10^6^ cells (lower and upper dashed lines, respectively). In the beginning of the starvation interval, we see an overshoot where the animal still grows although feeding has stopped. (**C**) A generic growth/degrowth dynamic is observed irrespective of the initial cell number, initial energy content or the feeding scheme. Any perturbation decays quickly and there is no strong dependence on feeding history. See *Figure 3 – figure supplement 1* for details.

**Figure 3 – figure supplement 3.**
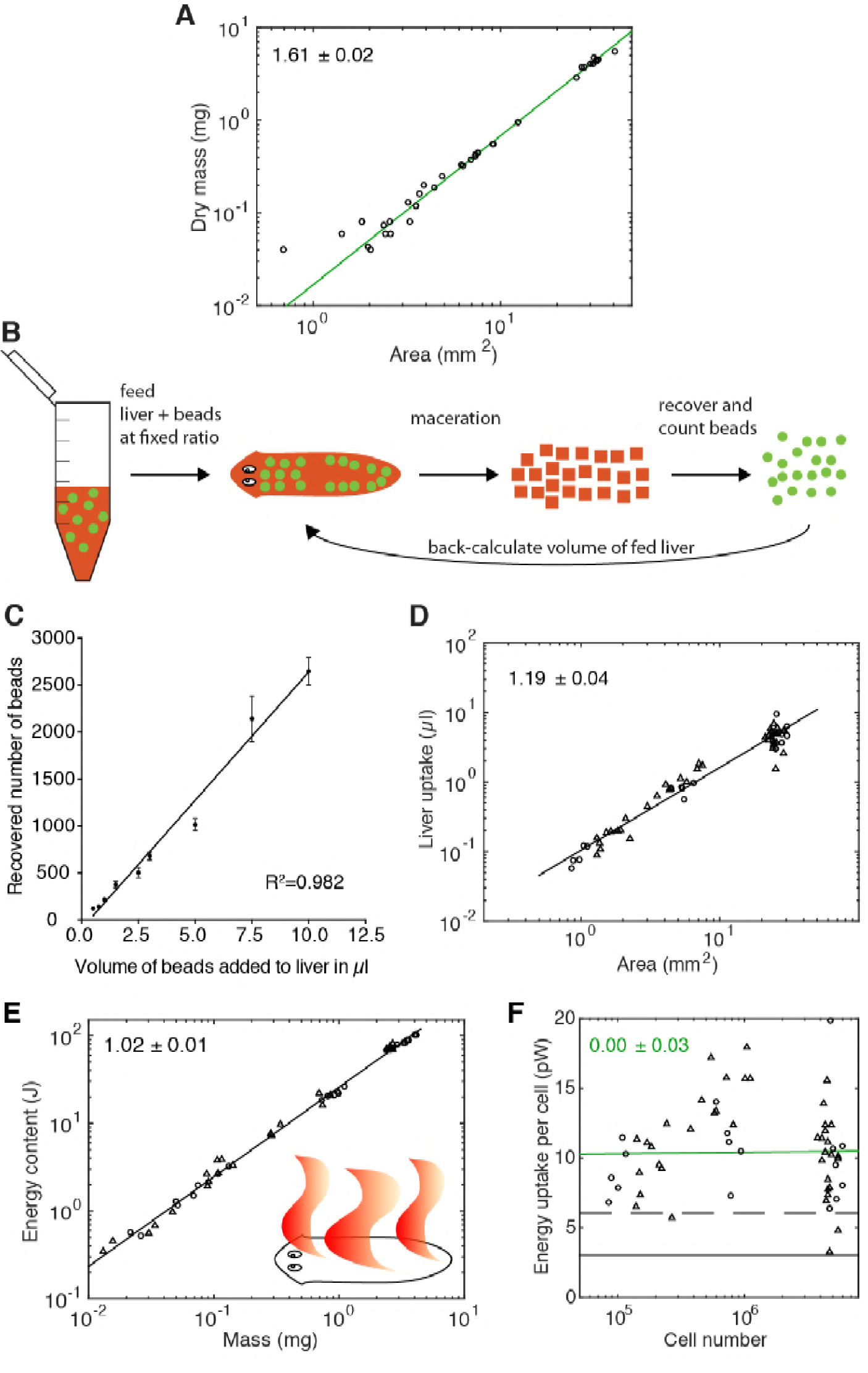
Paradigm validation. (**A**) Dry mass versus plan area scaling used to convert dry mass into cell numbers (via relationship in *Figure 2B*) (each data point = one individual animal, 2 independent experiments represented with 21 and 15 animals, respectively). The scaling exponent ± standard error was derived from a linear fit and represents the exponent *b* of the power law *y = ax*^*b*^. (**B**) Cartoon of food intake assay. A mixture of liver with a known concentration of spiked-in fluorescent beads as volume tracers is fed to the animals. Afterwards, individual animals are macerated and the number of recovered beads within a small volume fraction of the macerate are counted using a similar image-based approach as shown in *Figure 2A* (top). Extrapolation to the initial total sample volume reveals the total number of fed beads and finally, based on the known beads/liver ratio, the total volume of fed liver. (**C**) Recovery of a linear beads dilution series (R^2^= 0.982) fed to the animals demonstrates the validity of using beads as a volume tracer. The line represents a fitted linear regression to data from one experiment; error bars, standard deviation. (**D**) Volume liver taken in as a function of plan area. Volume and area represent the original measurements based on which we calculated energy intake using the known nutritional value of calf liver (“Nutrient report of calf liver,” 2016) and the density of human adult liver (Overmoyer et al., 1987) as well as organismal cell number using *Figure 2B*. Food intake is not dependent on feeding history, compare 2 weeks-(circles) with 3 weeks-starved (triangles) animals. The scaling exponent ± standard error was derived from a linear fit and represents the exponent *b* of the power law *y = ax*^*b*^. (**E**) Energy content (gross calorific value) as a function of dry mass. The energy content is not dependent on feeding history, compare 1 week-(circles) with 3 weeks-starved (triangles) animals. The scaling exponent ± standard error was derived from a linear fit and represents the exponent *b* of the power law *y = ax*^*b*^. See *Figure 3 – figure supplement 3 – source data 1* for data of (A), (D) and (E). (**F**) Food uptake per cell versus organismal cell number (*N*) (same data as in *Figure 3E*). Green line, fit to the data representing ingested feed energy. Scaling exponent ± standard error derived from linear fits and representing the exponent *b* of the power law *y = ax*^*b*^. The black solid line represents the physiological net energy uptake (~30 % of the ingested feed energy): i.e. the net amount of energy which is assimilated by the animal body following digestion in the gut (minus energy loss due to a potential elevated metabolism during feeding) and eventually fully metabolised, thus contributing to the measured heat production (predicted from *Figure 3D*, assuming a constant metabolic rate per cell of 1 pW). The conversion between gross and physiological energy (see *Figure 3F*) allows to predict the net ingested feed energy (dashed line) which is assimilated by the animal body (~60 % of ingested feed energy). This corresponds to a feed conversion ratio (ratio between feed mass and resulting gain in body mass, a measure for how efficient an organism converts feed into body mass) of ~2.6, similar to other aquatic animals (Tacon & Metian, 2008), see also discussion. Circles, 2 weeks-starved and triangles, 3 weeks-starved animals, indicate no obvious dependence on feeding history up to 3 weeks after feeding.

**Figure 4 – figure supplement 1.**
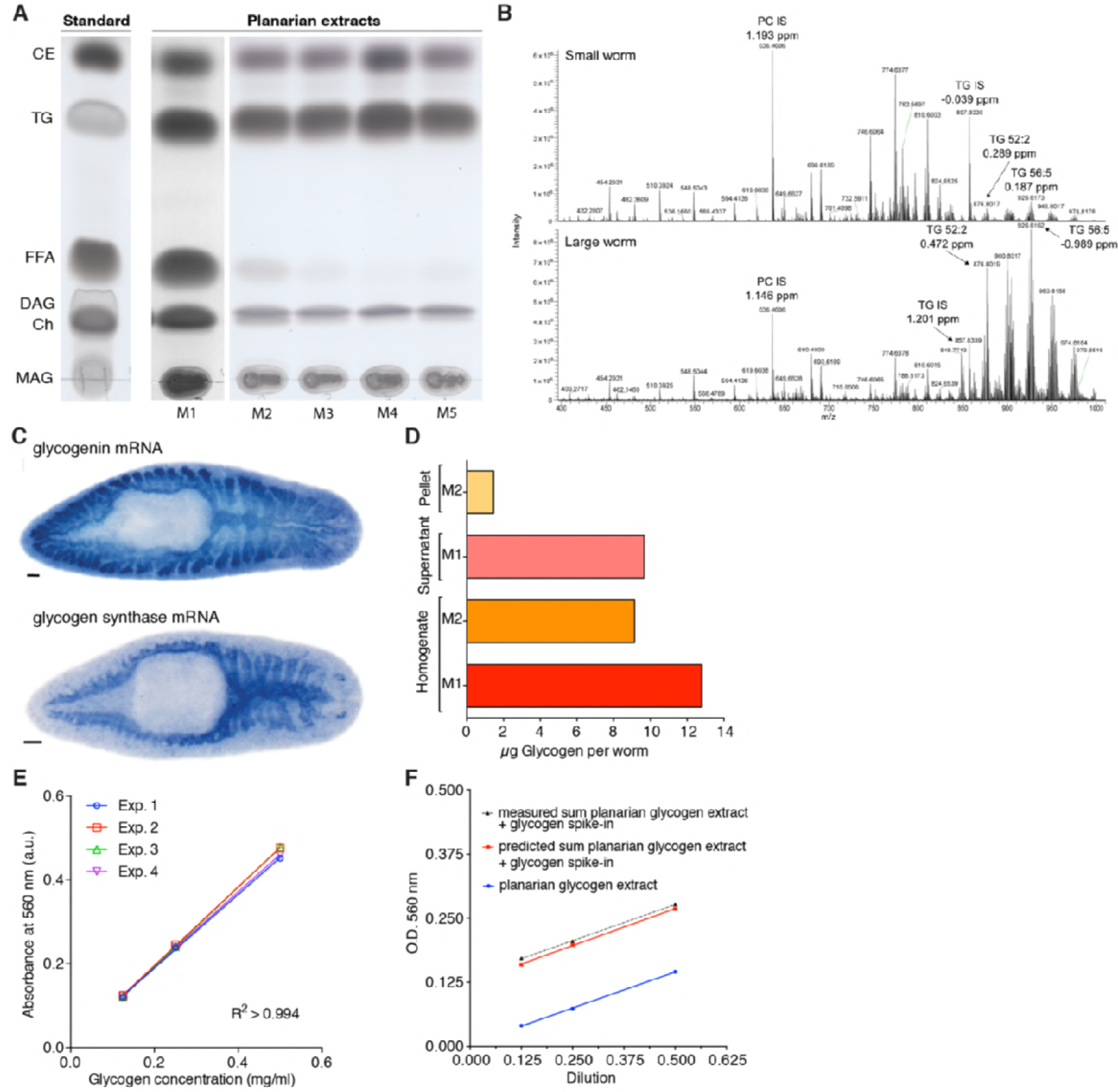
Alternative assays confirm lipid and glycogen storage in planarians. (**A**) Right, TLC (thin layer chromatography) analysis of planarian lipid extracts with different homogenization conditions to prevent triglyceride (TG) degradation: M1, Bligh and Dyer method (Bligh & Dyer, 1959); M2, 300 µl Isopropanol; M3, 1ml ice-cold Isopropanol; M4, ice-cold Isopropanol/Acetonitrile (1:1); M5, 1ml Isopropanol with 0.5% glacial acetic acid. The homogenization in ice-cold Isopropanol/Acetonitrile (1:1) generates the least amount of free fatty acids (FFAs), reflecting the least degradation rate of TGs. Left, TLC separation of the standards: CE, Cholesteryl linoleate; TG, Glyceryl trioleate; FFA, Linoleic acid; DAG, Dioleoylglycerol; Ch, Cholesterol; MAG, 1-Oleoyl-rac-glycerol. (**B**) Representative example of mass spectrum of small and large animals from positive mode (i.e. positively charged analytes). Highlighted are the internal standards for Phosphatidylcholine (PC IS) and Triglyceride (TG IS) and two of planarian endogenous Triglycerides (TG 52:2 and TG 56:5). The peak intensity of TG 52:2 and TG 56:5 is almost 14-fold higher in large worms compared to small worms while the intensity of PC IS and TG IS show only minor differences (~1.3-fold small vs. large). ppm, parts per million mass accuracy. See *Figure 4 – source data 2* for quantification of relevant lipid classes. (**C**) *in situ* hybridisation against glycogenin (left) and glycogen synthase mRNA (right) further supports our finding that planarian store sugar in the form of glycogen (see also *Figure 4C-D*). Scale bars, 100 µm. (**D**) Comparison of different glycogen extraction methods on total homogenate, supernatant and pellet fractions. M1: Water extraction, M2: hot alkali extraction. Water extraction and glycogen from the supernatant fraction were used in this study. (**E**) Glycogen standard curves from the glycogen content quantification assay. The lines represent linear regressions fitted to the data of each experiment. (n=4 independent experiments, 3 technical replicates). The line represents a fitted linear regression. (**F**) Linear dilution series of planarian glycogen extract (blue line) and the predicted cumulative dilution series of extract with spiked-in glycogen (concentration =0.16 µg/ul) (red line). The predicted sums for the extract with spiked-in glycogen (red line) are closely in line with the measured sums of extract and spiked-in glycogen (dotted grey line), demonstrating the linearity of the assay across the entire concentration range. The lines represent fitted linear regressions.

**Figure 5 – figure supplement 1.**
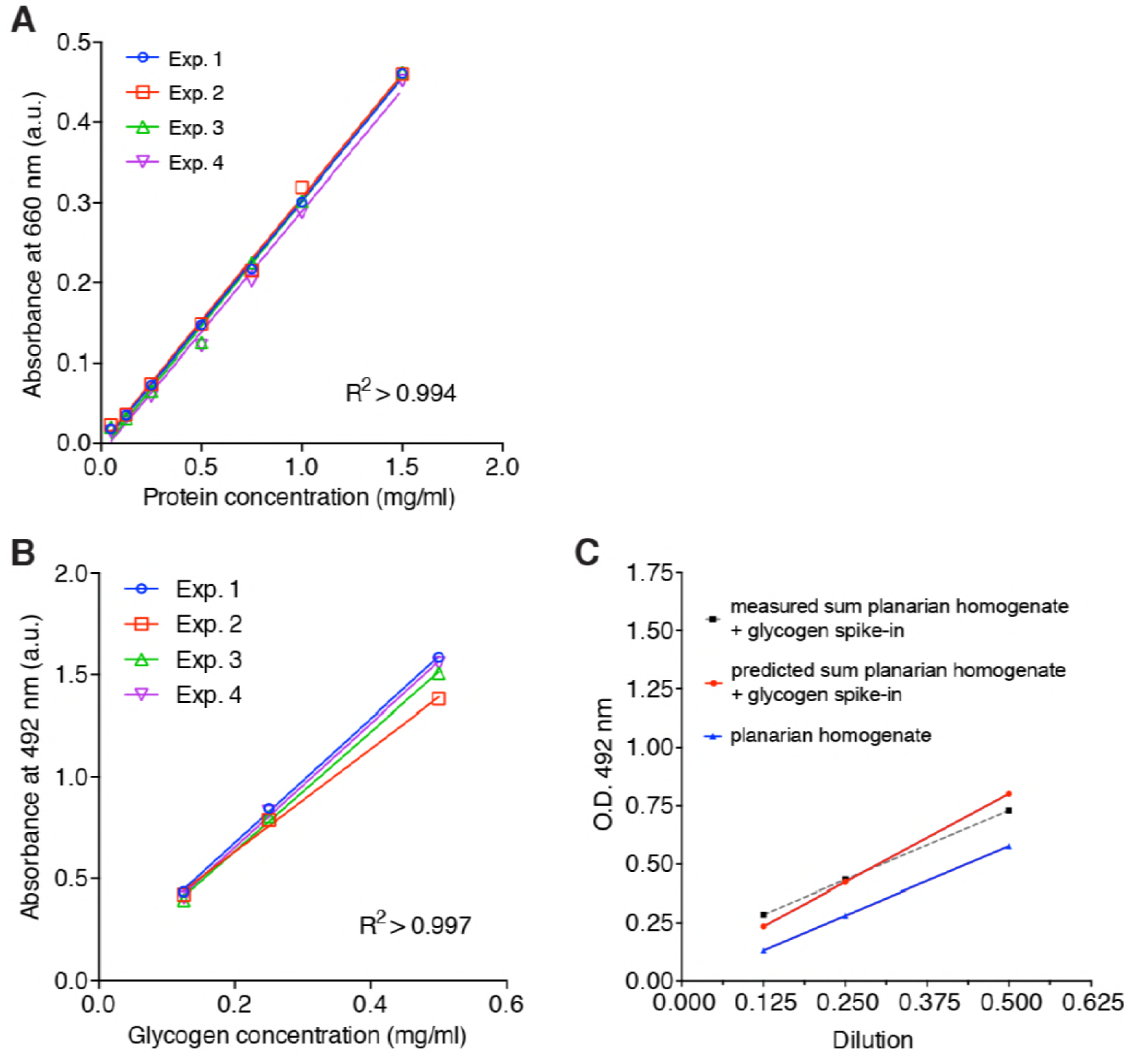
Standard curve protein and validation of total carbohydrate measurement. (**A**) Standard curves of the colorimetric protein quantification. The lines represent linear regressions fitted to the data of each experiment (n=4 independent experiments, 3 technical replicates). (**B**) Glycogen standard curves from the phenol-sulfuric acid method. The lines represent linear regressions fitted to the data of each experiment (n=4 independent experiments, 3 technical replicates). (**C**) Linear dilution series of planarian homogenate (blue line) and the predicted series of homogenate with spiked-in glycogen (conc.=0.25 µg/µl) (red line). The predicted sums for the homogenate with spiked-in glycogen (red line) are closely in line with the measured sums of homogenate and spiked-in glycogen (dotted grey line), demonstrating the linearity of the assay across the entire concentration range. The lines represent fitted linear regressions.

## List of figure supplements

Figure 1 – figure supplement 1.

Measurement of body size and metabolic rate.

Figure 2 – figure supplement 1.

Validation of image-based quantification of organismal cell number.

Figure 2 – figure supplement 2.

Degrowth rates are independent of feeding history.

Figure 3 – figure supplement 1 (text only).

Implementation of the theoretical model.

Figure 3 – figure supplement 2.

Further explanation of model paradigm 1.

Figure 3 – figure supplement 3.

Paradigm validation.

Figure 4 – figure supplement 1.

Alternative assays confirm lipid and glycogen storage in planarians.

Figure 5 – figure supplement 1.

Standard curve protein and validation of total carbohydrate measurement

## List of source data

Figure – source data 1.

Numerical data wet mass vs. dry mass measurements.

Figure 2 – source data 1.

Numerical data cell number measurements.

Figure 2 – source data 2.

Raw numerical data Histone H3 method (quantitative Western blotting)

Figure 2 – source data 3.

CellProfiler results tables image-based approach.

Figure 2 – source data 4.

MATLAB code for extraction of planarian body size.

Figure 2 – source data 5.

Numerical data growth/degrowth.

Figure 3 – source data 1.

Numerical data for Figure 3.

Figure 4 – source data 1

Raw images lipid droplet and glycogen.

Figure 4 – source data 2

Raw data lipid mass spectrometry, glycogen assay and statistics tables.

Figure 5 – source data 1

Raw data and statistics tables for measurement of other lipids, carbohydrates and protein.

Figure 1 – figure supplement 1 – source data 1

Raw data metabolic rate measurements.

Figure 2 – figure supplement 1 – source data 1

CellProfiler pipeline, numerical data, raw images and segmentation for validation of image-based cell counting.

Figure 3 – figure supplement 3 – source data 1

Numerical data for Figure 2 – figure supplement 3.

## List of supplementary files

List of scaling relationships.

